# Range-wide genetic population structure and environmental adaptation in the eastern oyster (*Crassostrea virginica*) provides insight for aquaculture

**DOI:** 10.64898/2026.03.30.715280

**Authors:** Madeline G. Eppley, Kiran Bajaj, Camille Rumberger, Elisabeth Leung, Nicole Mongillo, Jessica Small, Katie Lotterhos

## Abstract

Selective breeding in aquaculture is necessary to establish food security and meet demand for sustainably produced protein. An informed selective breeding program requires understanding how population structure, environmental adaptation, and human activities shape natural genetic variation in wild conspecifics. Unfortunately, wild variation remains poorly characterized for many commercially important aquaculture species. Here, we conduct the first range-wide study of genomic population structure for the eastern oyster (*Crassostrea virginica*) across thousands of miles (Texas, USA to Eastern Canada) using a 200K SNP array. We integrate population structure analyses, genotype-environmental associations, and structural variant detection to identify adaptive loci and quantify human-mediated genetic impacts. Our data confirms two ancestral clusters with a phylogeographic break between the Gulf and Atlantic (F_ST_ = 0.06) and highlights patterns of substructure within each region. We find evidence of unexpected patterns of genomic variation in two locations: evidence of Gulf ancestry in a mid-Atlantic estuary (Chesapeake Bay), and evidence of Atlantic ancestry in a Gulf estuary (Apalachicola Bay). While we cannot definitively determine the causes of these unexpected patterns, we show that they are consistent with direct and indirect human impacts in these estuaries. Genotype-environment association analyses with *in situ* temperature and salinity measurements were used to identify putatively adaptive loci, including SNPs within large structural variants (>1Mb). Our results identified genomic targets for aquaculture breeding programs aimed at climate resilience, reveal complex patterns of human impacts in managed systems, and demonstrate how seascape genomics can be used to improve aquaculture outcomes.

## Introduction

Expanding capacity in marine aquaculture presents new opportunities to address global food security. Realizing the full potential of aquaculture requires integrating knowledge of how natural variation, gene flow, and human management interact to shape the genomic landscape available for artificial selection. Rapid advances in genomic technologies have opened the door to improving aquaculture through selective breeding, but successful selection requires identifying specific genomic targets underlying economically important traits such as thermal tolerance, stress responses, and disease resistance (Andersen et al., 2025; Houston et al., 2020).

Unlike terrestrial agriculture, aquacultured marine species remain only a few decades removed from their wild progenitors and maintain substantial genetic diversity that may prove advantageous to adaptation in the face of changing environments (Diamond, 2002; Puritz et al., 2022). Shellfish species in particular present unique challenges for genomic selection. For instance, many molluscs have high levels of heterozygosity, which can contribute to high genetic load, larval mortality, and inbreeding depression (Plough, 2016). While genomic selection of aquacultured species has been effective for directly measurable traits like growth (*e.g.,* (Gutierrez et al., 2018; Marín-Nahuelpi et al., 2025), more complex traits such as salinity tolerance or disease resistance have remained a challenge to target (Hollenbeck & Johnston, 2018; Song et al., 2023).

Identifying adaptive loci and structural variants from wild populations is a priority for advancing genomic selection in aquaculture, particularly as climate change necessitates breeding for resilience to novel and variable environmental conditions (Dhillon et al., 2025). Marine species with broad geographic ranges frequently exhibit adaptive variation across environmental gradients in temperature, salinity, and disease pressure, so understanding the genomic basis of this local adaptation is essential for sourcing broodstock that will perform positively under expected environmental conditions (Grummer et al., 2019; Liggins et al., 2019). Structural variants, including chromosomal inversions, have emerged as important components of adaptive variation by preventing recombination and preserving locally adaptive gene complexes even under high gene flow (Hoffmann et al., 2004; Schaal et al., 2022; Tigano & Friesen, 2016; Wellenreuther & Bernatchez, 2018). Recent research in commercial and aquaculture species has shown that structural variants can be associated with local environmental adaptation (Hollenbeck et al., 2022) and economically important traits (Yang et al., 2024), yet these variants remain largely uncharacterized. Structural variants have also been shown to outperform SNPs alone in detecting finescale population structure in the American lobster (*Homarus americanus*), another important marine aquaculture species, suggesting it is necessary to investigate both structural variants and SNPs to understand population structure (Dorant et al., 2020).

In addition to identifying the genetic basis of adaptation, population structure is an important component of species management and can reveal ways that human activities have impacted the genetic composition of wild populations. For instance, human-mediated translocation of organisms has facilitated gene flow between naturally isolated populations, resulting in hybridization of domestic and wild individuals (Crispo et al., 2011). Human-mediated translocation of wild or cultured organisms can be deliberate (*e.g.,* stocking, assisted gene flow, restoration, hatchery operations) or accidental (*e.g.,* escapees, transport in ballast water), and is common across both terrestrial agriculture and aquaculture, altering populations at landscape scales. Human impacts on population structure have been observed in the European flat oyster in (Šegvić-Bubić et al., 2020), black lipped pearl oyster in (Lemer & Planes, 2012), and scallops in (Beaumont, 2000). Translocation between allopatric or genetically distinct populations can increase genetic diversity and resilience to environmental changes, but may also lead to loss of unique local genotypes, outbreeding depression, and maladaptation (Aitken & Whitlock, 2013; Edwards, 2015; Hornick & Plough, 2022; Zhao et al., 2024). Despite growing evidence for local adaptation in marine species (Akopyan et al., 2026; Conover et al., 2006; Lee et al., 2025; Sanford & Kelly, 2011; Whitlock & Gomulkiewicz, 2005; Xuereb et al., 2018; Yeaman & Otto, 2011), translocation across large scales for aquaculture and restoration continues to occur without a robust understanding of wild genetic structure. There is clear evidence of genetic impacts of admixture between local and transplanted individuals (Grant et al., 2017; McKay et al., 2005; Truskey et al., 2025; Varney et al., 2018), underscoring the need for careful genetic monitoring and management strategies to balance economic interests with conservation goals.

In addition to direct effects, human activities can also have indirect effects that shape genetic structure. Although climate change is the most widely studied (Bellard et al., 2012; Hoffmann & Sgrò, 2011), other human activities such as urbanization, water use, and overfishing can indirectly alter fitness landscapes (Allendorf & Hard, 2009; Pelletier & Coltman, 2018; Puritz & Toonen, 2011). For instance, changes to freshwater use far inland can have downstream effects on shellfish population health by altering the salinity in the estuary (Gray et al., 2009; Hintenlang et al., 2023; Petes et al., 2012; Pine et al., 2015). Ultimately, as both direct and indirect effects of human activities shape the selective environments that populations experience, understanding the interplay between natural genetic variation, environmental selection, and natural as well as human-mediated gene flow is essential for informed management of wild and cultured populations.

The eastern oyster *Crassostrea virginica* (Gmelin, 1791) is a heavily aquacultured shellfish species on the eastern coasts of the US and Canada with a complex history of human management. Pre-colonial wild oyster reefs were once sustainably managed by Indigenous fisheries (Reeder-Myers et al., 2022; Thompson et al., 2020), but contemporary overfishing has caused localized depletion (Beck et al., 2011; Jackson et al., 2001; Rothschild et al., 1994; Zu Ermgassen et al., 2012). To mitigate commercial losses, oysters were transported from neighboring regions to depleted areas (Bersoza Hernández et al., 2018; Kirby, 2004) as early as the 1920s (Churchill, 1920). Restoration efforts since the 1990s have revitalized the species economically and ecologically (Grabowski et al., 2012), yet source material for these efforts is often not genetically well characterized (Hornick & Plough, 2019). In particular, “local” hatchery spat does not necessarily represent wild local genotypes (Varney et al., 2018), and restoration efforts sometimes use hatchery-produced spat-on-shell, of which the genotypic source can vary (Bersoza Hernández et al., 2018). Investigating range-wide population structure will clarify the potential impacts of these management practices on wild populations and inform decision-making.

While existing research has provided important insight into the regional genetic structure of eastern oyster populations (Bernatchez et al., 2019; Puritz et al., 2022; Reeb & Avise, 1990; Thongda et al., 2018; Varney et al., 2009), no studies have comprehensively assessed population genomic structure of the species across its range. Eastern oyster populations span wide gradients in latitude, temperature (0° to 36°C), salinity (4 to 42 ppt), and disease pressure (Bernatchez et al., 2019; Carnegie et al., 2021; Carnegie & Burreson, 2011; Lowe et al., 2017; Sirovy et al., 2023). Developing an understanding of the genomic targets of selection in wild oyster populations to their local environment can inform breeding programs (Guo, 2009; Guo et al., 2023; Proestou et al., 2019), but such a comprehensive study has yet to be done.

To address these knowledge gaps, we use a high-density SNP array to sequence and analyze 746 eastern oyster individuals from 40 sites spanning the native range, producing the most comprehensive population genomic assessment to date. We investigate how genetic variation is partitioned across the native range, evaluate evidence for human-mediated gene flow and resulting introgression in regions of intensive management, and identify putative genomic targets of environmental selection. Our findings identify genomic targets for aquaculture improvement and generate insight for how human management practices shape the genetic landscape of wild populations.

## Methods

### Collection, DNA Extraction, and Genotyping

Eastern oyster samples were collected from 40 sampling locations across the species’ range along the Gulf coast of the United States and the Atlantic coasts of the United States and Canada (Supp. Table S1; Figure 1A). Collections targeted a sample size of 30 oysters per site and were sampled between May and October of 2022 and 2023. Oysters were collected by hand from intertidal zones, or by rake or dredge if the water depth at the site exceeded 5 feet at low tide. Collection efforts focused on sampling from established wild populations, avoiding proximity to aquaculture farms and recently restored sites. All samples were collected under state-level permits issued to the author(s) or collaborators who collected and shipped oysters.

**Figure 1.**
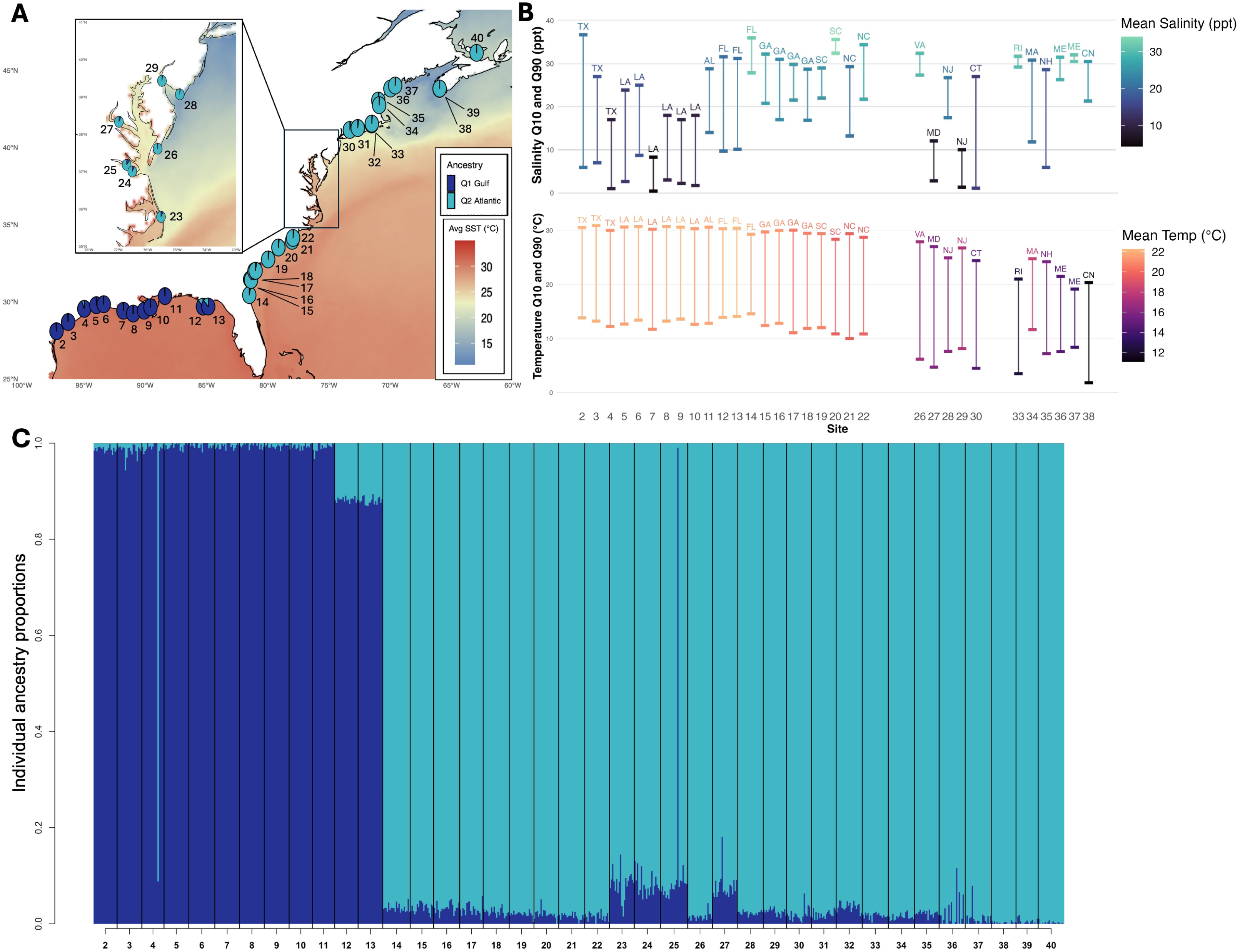
Environmental variation and ancestral groups across the seascape. A) Map of the seascape environment with the 40 sampling locations. Pie charts display averaged site-level estimates of Gulf (Q1, dark blue) or Atlantic (Q2, light blue) ancestry. B) Environmental variation across sampling sites showing Salinity_Q90_ and Salinity_Q10_ values (top), with bars colored by the average calculated salinity at each site, and Temperature_Q90_ and Temperature_Q10_ values (bottom). Environmental data was not available for some sites. C) Ancestry proportions for all numbered sites (2-40 from Figure 1) from sNMF analysis. Q1 (dark blue) vertical bars correspond with the Gulf ancestral cluster, and Q2 (light blue) vertical bars correspond with the Atlantic ancestral cluster.

Collected oysters were then shipped live on ice to the Northeastern University Marine Science Center in Nahant, MA. Oysters were kept alive in a 5℃ fridge for up to 72 hours until the oysters were shucked and tissue biopsies of gill were taken. Biopsies were preserved in 95% molecular grade ethanol and stored at -80℃.

Genomic DNA was extracted from 18-22 mg of gill tissue using an optimized Qiagen DNeasy Blood and Tissue kit protocol to yield high molecular weight DNA. Modifications included extending the pre-elution incubation step to 10 minutes at room temperature and eluting in low TE buffer. DNA quantity and quality were assessed with a PicoGreen Assay and 2% gel electrophoresis. Samples meeting minimum thresholds (>20 ng/uL with distinct high molecular weight bands >3kb and minimal streaking) were concentrated to 400 ng in 30 uL molecular-grade water, dried, and shipped for genotyping. Of the 1,120 samples collected, 800 samples from all 40 populations (*n* = 20 / population) were randomized across 10 plates and genotyped using a Affymetrix Axiom myDesign Custom Array with 200K SNPs at Neogen Genomics (Lincoln, NE). The 200K array was based on a 600K array developed from variant calls aligned to the haplotig-masked *Crassostrea virginica* reference genome (C_virginica-3.0, GenBank assembly GCA_002022765.4) (Gómez-Chiarri et al., 2015; Guo et al., 2023; Modak et al., 2021; Puritz et al., 2024).

### SNP Filtering

Genotype calls were made globally by Neogen Genomics after completion of all plate runs. After quality control and removing technical replicates, 745 individuals were retained for analysis. Raw VCF files were processed in R v4.2.2 (R Core Development Team, 2021) into a diploid genotype matrix using extract.gt() in vcfR (Knaus & Grünwald, 2017). The SNP array contained 195,025 high-density markers at nuclear and mitochondrial loci.

We first filtered for SNPs with >10% missing data across individuals (*n* = 5,756 SNPs removed), followed by minor allele frequency (MAF) filtering to retain SNPs with MAF > 0.05, resulting in 153,921 retained loci. Missing genotypes were imputed using the snmf() and impute() functions from the package LEA v3.6.0 (Frichot & François, 2015). We ran snmf() for K=1 through K=5 ancestral populations with 10 repetitions each, and K=2 was selected based on the lowest cross-entropy value. Using the best K=2 run, missing genotypes were imputed with mode-based imputation, which assigns the most likely genotype at the missing position as determined by the genotype frequency estimate at that site within the ancestral group. This generated complete data across all individuals at 153,921 loci, hereafter referred to as the *global full SNP dataset (Table 1)*.

**Table 1.** Summary of filtered SNP datasets used in downstream analyses. Regional datasets were re-filtered for minor allele frequency within each region after subsetting from the global dataset.

| Dataset | Number of individuals | Number of SNPs | Analyses |
| --- | --- | --- | --- |
| *Global full SNPs* | 745 | 153,921 | pcadapt, outFLANK, global lfmm2 temperature, local PCA, introgression |
| *Global thinned SNPs* |  | 115,171 | Global PCA, global sNMF, pcadapt (parameterization), outFLANK (parameterization), global lfmm2 temperature (parameterization) |
| *Gulf full SNPs* | 220 | 114,465 | Gulf lfmm2 salinity (test) |
| *Gulf thinned SNPs* |  | 89,641 | Gulf PCA, Gulf sNMF, Gulf lfmm2 salinity (parameterization) |
| *Atlantic full SNPs* | 523 | 130,597 | Atlantic lfmm2 salinity (test) |
| *Atlantic thinned SNPs* |  | 101,847 | Atlantic PCA, Atlantic sNMF, Atlantic lfmm2 salinity (parameterization) |

Next, we created a *global thinned SNP dataset* (Table 1) for estimating population structure and for neutral parameterization in genome scans. The global thinned SNP dataset was generated by removing SNPs in linkage disequilibrium from the full SNP matrix using the package bigsnpr v1.11.3 (Privé et al., 2018), which applies Single Value Decomposition (SVD) to reduce correlated SNPs over multiple iterations. The full SNP matrix was converted into bigSNP format, then thinned with snp_autoSVD()with default parameters (thr.r2 = 0.2; 500 bp window). After convergence, 115,171 SNPs were retained in the global thinned SNP matrix which was used for all population structure analyses (Table 1).

As Gulf and Atlantic populations are divergent, we subset the global full and thinned SNP matrices by region and re-applied MAF > 0.05 filtering within each to generate Gulf and Atlantic SNP datasets (Table 1).

### Population Structure

#### Principal Components Analysis

Population genetic structure was examined with principal components analysis (PCA) using the global thinned SNP dataset from snp_autoSVD() for all individuals (*n* = 745). Separate PCAs were also conducted on the Gulf (*n* = 220, *Gulf thinned SNP dataset*) and Atlantic (*n* = 523, *Atlantic thinned SNP dataset*) using prcomp() from the package stats v4.2.2. The percentage of variance explained by each PC axis was calculated as the squared eigenvalue divided by the sum of all squared eigenvalues, multiplied by 100.

#### Ancestry

Population ancestry coefficients were inferred on the global thinned SNP dataset, the Gulf thinned SNP dataset, and the Atlantic thinned SNP dataset using sparse non-negative matrix factorization (sNMF) implemented in LEA (Frichot & François, 2015). For each dataset, we tested *K* values between 1-20, and identified the lowest cross-entropy value. Individual ancestry coefficients were extracted from the Q-matrix for data visualization. For visualization on maps, the mean ancestry coefficient for each site was calculated by averaging individual ancestry proportions at that sampling location.

#### Population genetic statistics

Pairwise F_ST_ among all sampling locations were calculated using the *global thinned SNP dataset* using the Weir and Cockerham method (Weir & Cockerham, 1984) with the package hierfstat v0.5.11 (Goudet & Jombart, 2024). Observed heterozygosity (H_o_), expected heterozygosity (H_e_), and inbreeding coefficient (F_IS_) were calculated for each population using the basic.stats() function in hierfstat (Goudet & Jombart, 2024) on the *global thinned SNP matrix*.

### Local PCA

To characterize islands of divergence across the genome we performed local PCA in sliding windows of 100 SNPs with a step size of 20 SNPs for each chromosome on the *global full SNP dataset* using the locstruct v.0.0.0.9 package (Li & Ralph, 2019). We calculated pairwise distances between windows based on PC coordinates, and visualized these distances using multidimensional scaling (MDS) plots colored by window position. We plotted MDS1 and MDS2 coordinates against genomic position (Mbp) to identify regions of the genome with “outlier” ancestry following (Li & Ralph, 2019). To determine if these “outlier” regions represented large structural variants (e.g., inversions), we then performed standard PCA on each “outlier” region-ordered samples by PC1 loadings in a heatmap to visualize haplotype clustering patterns.

### Outliers for genetic differentiation

#### Pcadapt outlier analysis

To identify loci that show genetic differentiation above that expected by population structure we first conducted a principal component-based outlier detection analysis using the package pcadapt v4.3.5 (Luu et al., 2017). We parameterized pcadapt analyses with the *global thinned SNP dataset* and ran the analysis on the *global full SNP dataset* with *K*=2 (see Population Structure - Ancestry) groups to correct for structure as recommended by Lotterhos 2019 (Lotterhos, 2019). To assess statistical significance of detected outlier SNPs, a Bonferroni correction with α = 0.1 was applied to control for multiple tests.

#### OutFLANK outlier analysis

Neutral parameters were estimated on the global thinned SNP matrix (left and right trim fractions = 0.05; minimum heterozygosity = 0.1). Outlier detection was then performed on the global full SNP matrix using a *q*-value threshold of 0.01 (Lotterhos & Whitlock, 2014).

### Environmental data and genotype-environment association analysis

Temperature and salinity data for each sampling location (Supp. Table S2) was downloaded from sources with routine monitoring (*e.g.,* National Estuarine Research Reserve System, National Parks Service) (Helmuth et al., 2006; Nadeau et al., 2017). Raw environmental data were converted into a standardized format using a custom R script (Data Accessibility).

Sampling locations were retained in genetic-environment association analysis if their data were high-quality by meeting the following minimum requirements: 1) at least 1 year of measurements, 2) a minimum of 6 months of data per year, with at least 1 month capturing seasonal highs and lows, 3) at least 1 measurement per day, and 4) recent data collected within the 5 years prior to sampling. Out of the 40 sampling locations, 32 locations met these quality control measures. Extreme values were filtered out to remove data errors (valid ranges: 0-40 ppt for salinity, 0-42℃ for temperature). For each sampling location, we compiled the mean, maximum, and minimum annual values, standard deviation, and quantiles (10th - low and 90th - high percentiles) for salinity and temperature, hereafter referred to as Salinity_Q90,_ Salinity_Q10,_ Temperature_Q90,_ and Temperature_Q10_. Variables with significant pairwise correlations (*r* > 0.8) were removed. Quantiles and standard deviations were used in downstream analysis.

To identify loci putatively under environmental selection, we conducted latent factor mixed model analyses (LFMM) in the LEA package (Frichot & François, 2015). Temperature analyses were run globally across the 32 sampling locations for which we had high-quality environmental data (609 individuals, *K* = 2), as Gulf sampling locations exhibited minimal temperature variation. Salinity analyses were run separately for the Atlantic (387 individuals, *K* = 5) and Gulf (220 individuals, *K* = 2) as salinity regimes vary across each region. For each analysis, we parameterized the model with lfmm2() on the corresponding thinned SNP matrix (Table 1) and tested for associations on the corresponding full SNP matrix with lfmm2.test() using a Fisher’s test to obtain a significance value for each locus. We applied Bonferroni correction with α = 0.1 to identify outlier SNPs.

### Gene Ontology Analysis

We performed gene ontology (GO) enrichment analysis on outlier loci identified through outFLANK, lfmm2, and pcadapt with ShinyGO v0.85.1. Outliers were mapped to gene regions using the *C. virginica* genome annotation from Ensembl Metazoa (release 62) (Yates et al., 2022). The package rtracklayer v1.58.0 (Lawrence et al., 2009) was used to import the GFF3 file and GenomicRanges v1.50.2 was used to identify connections between outlier SNPs and annotated genes. Gene identifiers were extracted and used to query Ensembl Metazoa with the biomaRt package v2.54.1 (Durinck et al., 2009) to retrieve gene descriptions and GO terms. For functional enrichment analysis, we used ShinyGO v0.82 with the *C. virginica* reference (C_virginica-3.0, GenBank assembly GCA_002022765.4). We considered terms with FDR < 0.05 as significantly enriched.

### Analysis of genomic regions exhibiting unusual patterns of ancestry

For sites in two large estuaries with unexpected patterns of ancestry which resembled introgression from a genetically distinct population (Apalachicola Bay - sites 12, 13; Chesapeake Bay/Pamlico Sound - sites 23, 24, 25, 27), we conducted further analyses. We used a permutation analysis based on diagnostic loci to develop a deeper understanding of the genomic regions showing introgression from a distinct genetic population.

First, we identified highly differentiated loci that could distinguish between Gulf and Atlantic populations. We removed sampling locations with unusual patterns of ancestry (*n* = 6) and split the remaining 33 sampling locations into “Atlantic” (23 sampling locations; *n* = 444 individuals, excluding sampling locations with an average proportion of Gulf ancestry (Q1) ≥ 0.02) and “Gulf” (10 sampling locations; *n* = 185 individuals, excluding sampling locations with an average proportion of Atlantic ancestry (Q2) ≥ 0.01) populations. We calculated F_ST_ between the “Atlantic” and “Gulf” populations using the MakeDiploidFSTMat()function in OutFLANK (Whitlock & Lotterhos, 2015). We then filtered for highly differentiated loci (F_ST_ > 0.8) between regions, loci that we define as *diagnostic loci*. As the reference genome of the eastern oyster was sequenced from a selectively-bred line developed from Delaware Bay (Atlantic) ancestry (Puritz et al., 2024), the vast majority of reference (REF) alleles had high frequency in the Atlantic and low frequency in the Gulf, and we subset our analysis to these loci for ease of interpretation (5,542 loci).

To determine the genomic regions that showed unusual patterns of ancestry, we divided all sampling locations into one of four groups: *Atlantic* (described above)*, Gulf* (described above), *Chesapeake* (4 sampling locations including Pamlico Sound, an Atlantic estuary showing Gulf introgression), or *Apalachicola* (2 sampling locations, a Gulf estuary showing Atlantic introgression). At each diagnostic locus, we calculated the frequency of the reference (REF) allele in the Gulf, Atlantic, Apalachicola, and Chesapeake. Then, for each diagnostic locus we calculated the allele frequency change (!”) at the REF allele between the focal group showing mixed ancestry and the ancestral group as Δp = p_FOCAL_ - p_ANCESTOR_. The null hypothesis is that there is no allele frequency change (H_0_: Δp = 0). In the *Atlantic* where REF alleles are high in frequency, introgressing alleles would be in lower frequency in the *Chesapeake*, Δp = p_CHESAPEAKE_ - p_ATLANTIC_ and negative values of Δp indicate allele frequency change in the *Chesapeake* toward the *Gulf* (H_A_: observed Δp ≤ null Δp). In the *Gulf* where REF alleles are low in frequency, introgressing alleles would be in higher frequency in *Apalachicola*, and Δp = p_APALACHICOLA_ - p_GULF_ and positive values of Δp indicate allele frequency change in *Apalachicola* toward the *Atlantic* (H_A_: observed Δp ≥ null Δp). To assess statistical significance of the observed allele frequency changes, we then used permutation. For each comparison at each diagnostic locus, we shuffled population labels without replacement (*n* = 1000) and recalculated Δp. The permuted distribution at that locus represents a null distribution of allele frequency changes between the two groups. We then calculated the empirical p-value for the observed Δp at that locus as (r+1)/(n+1), where *n* is the number of permutations and r is the number of these replicates that produce a Δp more extreme than that calculated for the actual data in the direction of the alternative hypothesis (North et al., 2002). To correct for multiple testing, we applied Benjamini-Hochberg false discovery rate (FDR) correction at α = 0.05. Thus, loci with significant Δp in the direction of the alternative hypothesis showed patterns of ancestry that were significantly different from that expected by random. Loci with significant Δp were annotated and GO terms for associated genes were retrieved to identify functional consequences of introgression.

## Results

### Genotyping

After passing facility quality control, 745 samples were retained for further analysis (93% pass rate). All but one individual from the Laguna Madre, TX sampling site (Pop. 1, *Supp. Table 1*) failed genotyping, so this population was removed from further analysis.

### Population Structure

The ancestry analysis on the global thinned SNP dataset (*n* = 745 individuals) with two genetic clusters (*K* = 2) separated the Gulf and Atlantic ancestral groups (Figures 1A, 2E, Supp. Fig. S1). Principal component analysis separated Atlantic and Gulf individuals along PC1 (31% variance explained), and PC2 separated northern and southern Atlantic individuals (13.4% variance explained) (Figure 2A). Between Gulf and Atlantic populations, genome-wide pairwise *F_ST_* reached a maximum of 0.0647 between Calcasieu Lake, Louisiana (site 6) and Prince Edward Island, Canada (site 40) (Figure 2C). Inbreeding (*F_IS_*) was negative across all sampling locations, ranging from -0.285 (Calcasieu Lake, LA) to -0.222 (Long Island Sound, CT) (Table S1). Gulf sampling locations had a more negative average *F_IS_* (-0.283) than Atlantic sampling locations (-0.231), consistent with higher observed heterozygosity in the Gulf (Table S1).

**Figure 2.**
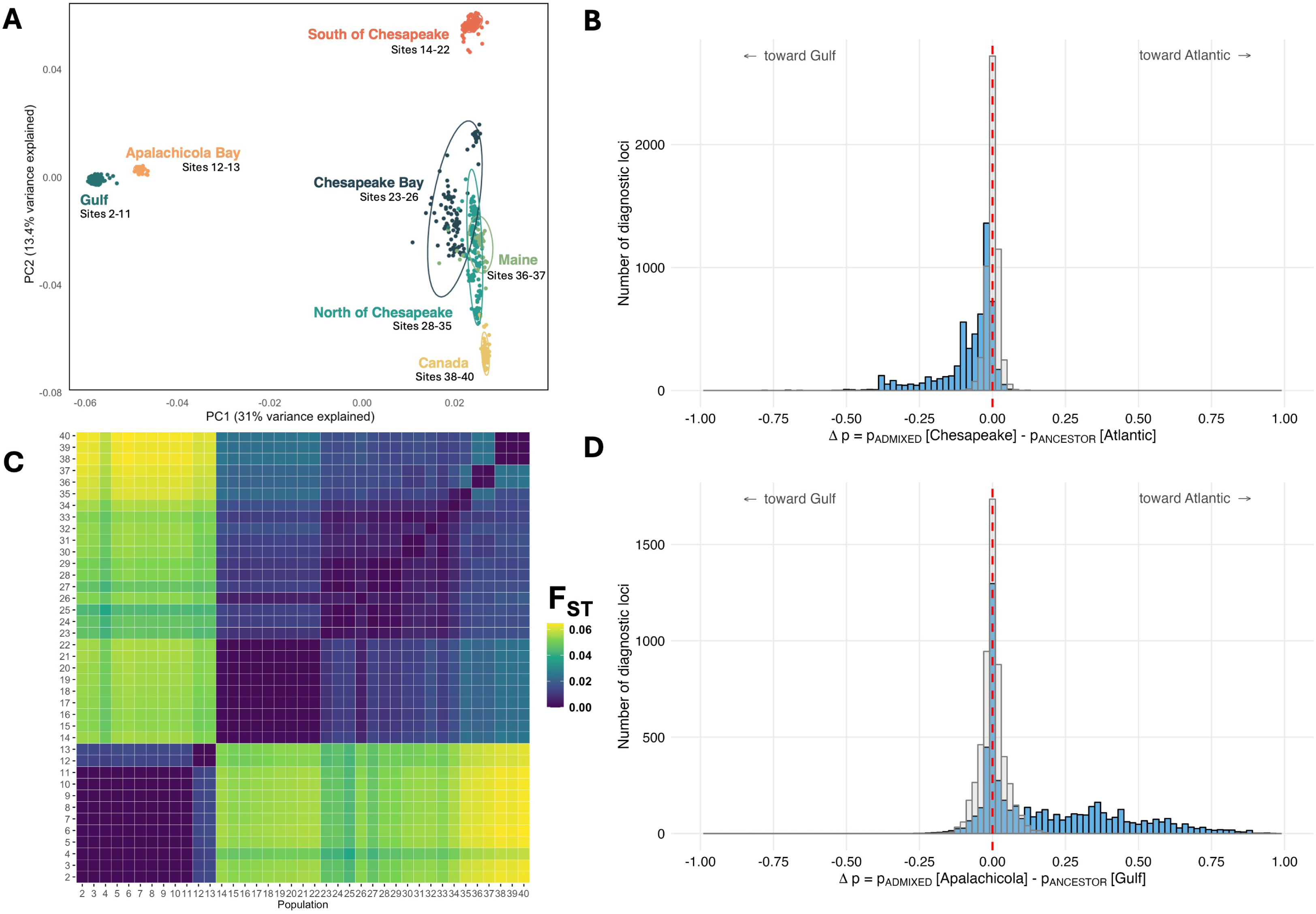
Population structure of eastern oysters across the seascape. A) Principal component analysis of 115k SNPs from 745 individuals from Gulf (sites 2-11) and Atlantic (sites 14-40) populations, separated along PC1 (31% variance explained) and PC2 (13.4% variance explained). B) Distribution of *Δp* values for Atlantic-vs.-Gulf diagnostic loci in the Chesapeake and Pamlico Sound, NC. Where *Δp* values are negative, allele frequency is changing towards Gulf composition. C) Heatmap of pairwise genetic differentiation (F_ST_) between all sampling sites. Lighter colors indicate higher genetic differentiation (maximum global F_ST_ = 0.0647) and darker colors indicate genetic similarity. D) Distribution of *Δp* values for Atlantic-vs.-Gulf diagnostic loci in Apalachicola Bay. Where *Δp* values are positive, allele frequency is changing towards Atlantic composition.

PCA and sNMF analyses on the Gulf and Atlantic subpopulations revealed finer-scale structure. In the Gulf, highest support was found for *K* = 2 (Supp. Fig. S2), which separated Apalachicola Bay from the rest of the Gulf, where PC1 explained 40.2% of variance and PC2 explained 1.4% of variance (Figure 3). Within the Gulf sites, maximum genome-wide pairwise *F_ST_* = 0.015 (Figure 2C). In the Atlantic, the cross entropy declined with *K*, an expected value associated with isolation-by-distance (Supp. Fig. S3). We chose to present *K* = 5, which revealed a Southern Atlantic cluster (Florida to North Carolina), three distinct northern clusters (in New Hampshire, Maine, and Canada), with complex patterns of ancestry in the mid-Atlantic (Figure 4). PC1 explained 42.3% of variance and PC2 explained 1.6% of variance (Figure 4). Within the Atlantic sites, maximum pairwise *F_ST_* = 0.0258 (Figure 1C).

**Figure 3.**
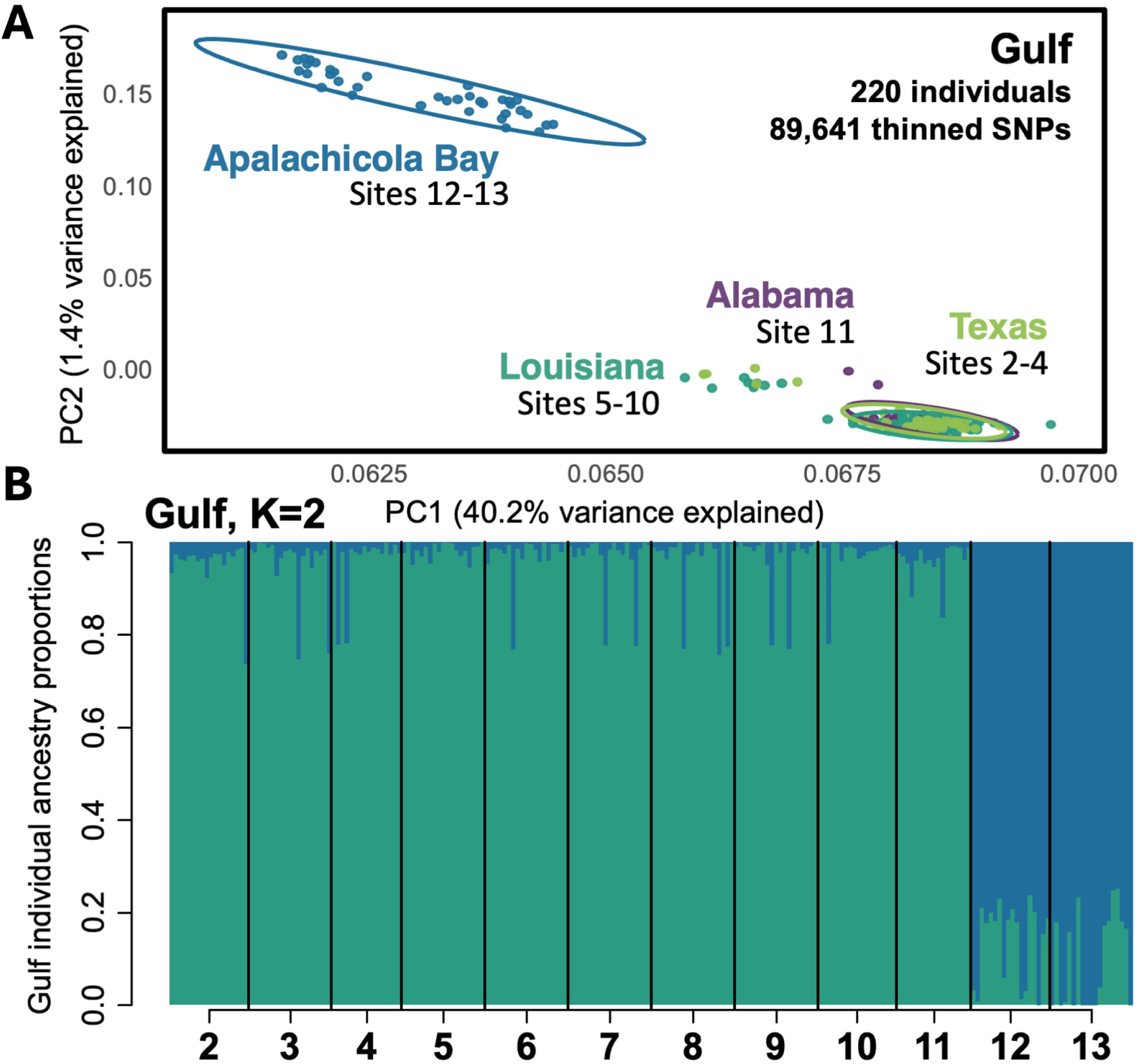
Regional population structure along the Gulf coast. Principal components analysis of Gulf populations (sites 2-13) and sNMF ancestry proportions with highest-support (*K* = 2) for Gulf individuals ordered by sampling site (site numbers correspond to map in Figure 1).

**Figure 4.**
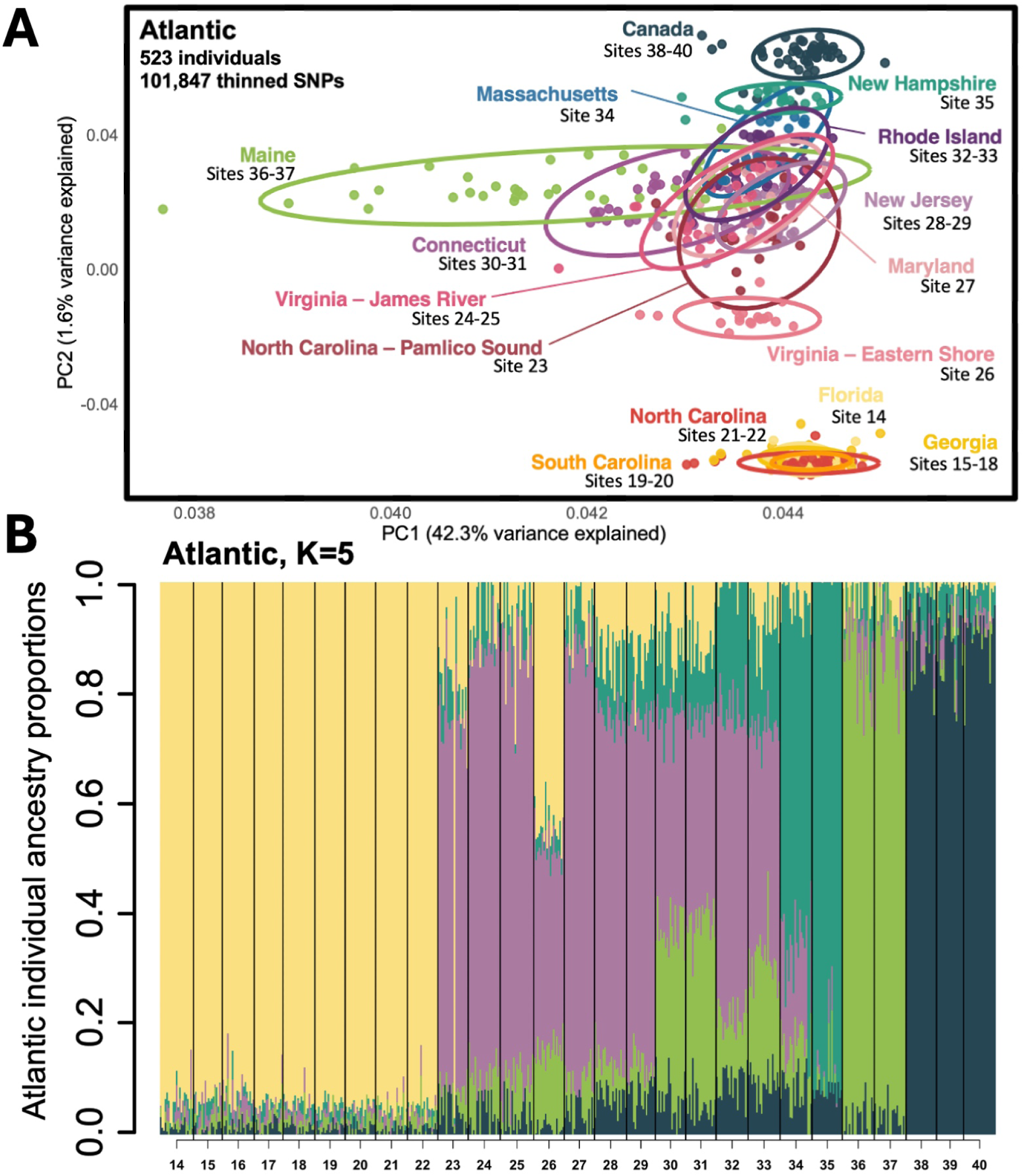
Regional population structure along the Atlantic coast. Principal component analysis of Atlantic populations (sites 14-40) and sNMF ancestry proportions with highest-support (*K* = 5) for Atlantic individuals ordered by sampling site (site numbers correspond to map in Figure 1).

### Analysis of estuaries exhibiting unusual patterns of ancestry

Of the 153,921 SNPs tested in the global full dataset, 5,542 (3.6%) were identified as diagnostic loci (*F_ST_* > 0.8) between Gulf and Atlantic populations, distributed across the genome in all 10 chromosomes. In the Chesapeake Bay group, the distribution of *Δp* values was mostly negative (as extreme as *Δp =* -0.8) (Figure 2B), indicating that Chesapeake allele frequencies have shifted away from the Atlantic ancestral population towards Gulf origin alleles. In Apalachicola Bay, the distribution of *Δp* values was mostly positive (as extreme as *Δp =* 1.00) (Figure 2D), indicating that Atlantic-origin alleles are increasing in frequency. Using the permutation-based approach, we tested if the allele frequencies at these diagnostic loci in two admixed populations (Chesapeake Bay; Apalachicola Bay) were significantly different from ancestral source populations (Atlantic; Gulf). In the Chesapeake Bay, a large proportion of loci were below the lower tail of the null distribution (Figure 2B), and after FDR correction (<0.05), 3,073 loci had significant negative *Δp* values (Figure 8A). In Apalachicolca Bay, many loci were above the upper tail of the distribution (Figure 2D), and after FDR correction (<0.05), 2,729 loci had significant positive *Δp* values (Figure 8B). Across the genome, diagnostic loci were broadly distributed, and many exceeded permutation confidence intervals (Figure 8A-B).

**Table 2.**
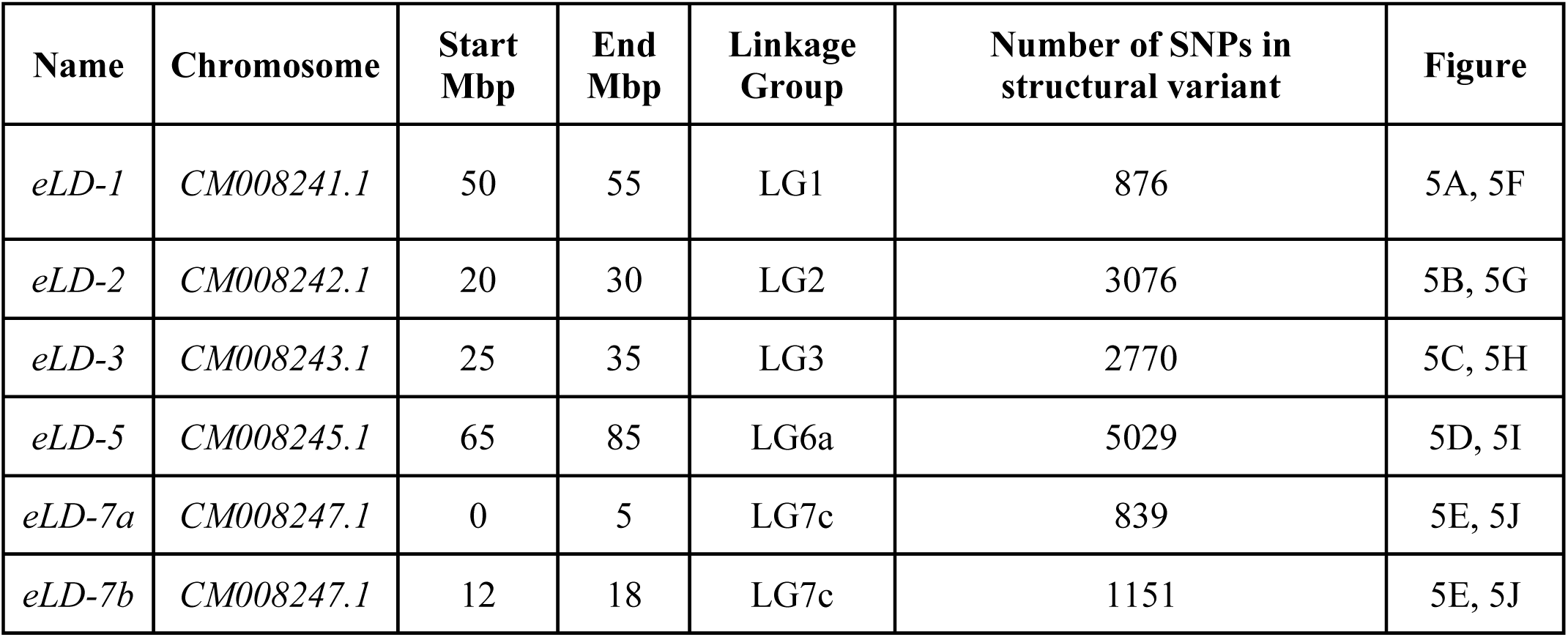
Location of detected structural variants and islands of divergence across the genome. Abbreviations: eLD = extended Linkage Disequilibrium region, Mbp = megabase pairs; SNPs = single nucleotide polymorphisms. Linkage groups (LGs) represent sets of loci that are inherited together by recombination patterns which may not align with physical chromosomes due to genome misassemblies (Supp. Fig. S5) or gaps in reference assembly.

| Name | Chromosome | Start Mbp | End Mbp | Linkage Group | Number of SNPs in structural variant | Figure |
| --- | --- | --- | --- | --- | --- | --- |
| *eLD-1* | *CM008241.1* | 50 | 55 | LG1 | 876 | 5A, 5F |
| *eLD-2* | *CM008242.1* | 20 | 30 | LG2 | 3076 | 5B, 5G |
| *eLD-3* | *CM008243.1* | 25 | 35 | LG3 | 2770 | 5C, 5H |
| *eLD-5* | *CM008245.1* | 65 | 85 | LG6a | 5029 | 5D, 5I |
| *eLD-7a* | *CM008247.1* | 0 | 5 | LG7c | 839 | 5E, 5J |
| *eLD-7b* | *CM008247.1* | 12 | 18 | LG7c | 1151 | 5E, 5J |

### Distinct patterns of local ancestry on chromosomes

Local PCA analysis revealed 6 genomic regions with distinct patterns of local ancestry on five of the ten chromosomes (Figure 5, Supp. Fig. S4). These patterns were consistent with extended patterns of linkage disequilibrium that local PCA detects, but did not show the characteristic pattern of three distinct genetic clusters in PC space expected by inversions (Kess et al., 2021; Li & Ralph, 2019; Lotterhos, 2019), so we refer to them as “extended LD regions” here. PC analysis did reveal, however, that multilocus genotype clusters tended to separate by major geographic region (Figure 5, Gulf in cool colors vs. Atlantic in warm colors). In total, 13,741 SNPs fell within extended LD regions across the genome (Table 2).

**Figure 5.**
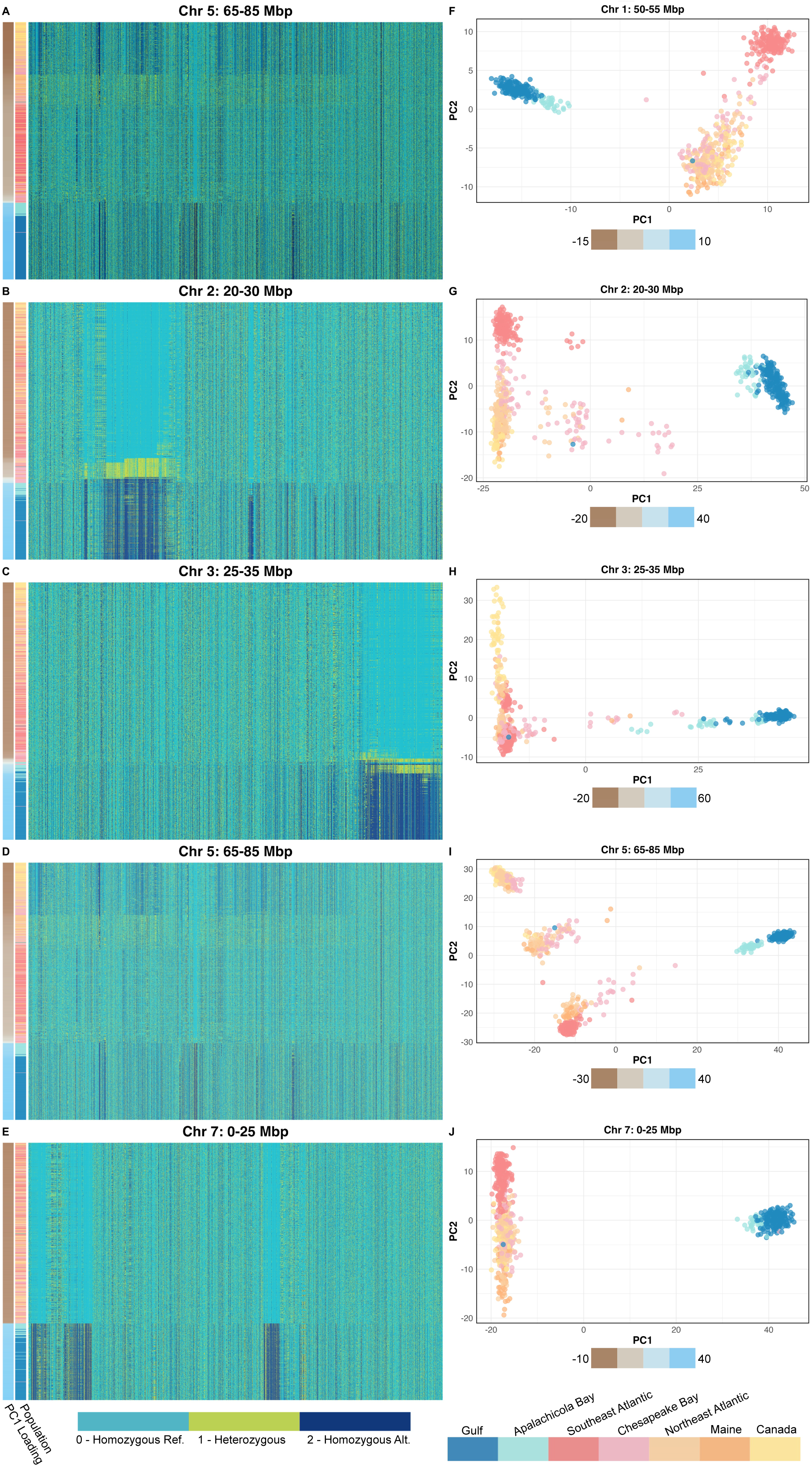
Regions of extended linkage disequilibrium across the genome. Figure 5A-E shows genotype heatmaps for windows of extended linkage disequilibrium on chromosomes 1, 2, 3, 5, and 7 (Table 2). Rows represent individuals ordered by PC1, and columns contain SNPs within each window. Genotypes are colored as homozygous reference (0, light blue), heterozygous (1, lime green) and homozygous alternate (2, dark blue). Figures F-J show the corresponding PCA on the same window, with points colored by Population. Each heatmap and PCA pair highlights the pattern of extended linkage disequilibrium within detected regions.

On chromosomes 1 (50-55 Mbp), 2 (20-30 Mbp), 3 (25-35 Mbp), and 7 (7a: 0-5Mbp, 7b: 12-18 Mbp), genotype heatmaps revealed inversion-like patterns (Figure 5), but the PCA patterns were more complex (Supp. Fig. S4). The largest extended LD region was detected on chromosome 5 (65-85 Mbp), which showed three distinct PCA clusters in the Atlantic and an additional two clusters in the Gulf.

### Environmental Selection

#### Temperature

Genotype-environment association analysis (lfmm2) performed on the global dataset of all individuals (*n* = 745) identified 1318 outlier loci corresponding to 388 annotated genes associated with high temperature (Temperature_Q90_) (Figure 6A) and 734 outlier loci corresponding to 201 annotated genes associated with low temperature (Temperature_Q10_) (Figure 6B). From the high temperature analysis, a strong signal of differentiation was observed within an extended LD region on chromosome 2 (20-30 Mbp, LG2, Table 2), where 8 outlier SNPs were detected within *Myosin heavy-chain muscle-like* (LOC111120907, max −log_10_(*P*) = 47.55), a gene involved with heat shock protein *HSP90*. On chromosome 3, outlier loci were detected within *Advillin-like* (LOC111122703, max −log_10_(*P*) = 13.26), a gene involved in cytoskeletal dynamics. Additionally, outlier loci on chromosome 3 were identified within *DnaJ homolog subfamily C member 7-like* (LOC111122700, max −log_10_(*P*) = 18), a heat shock protein in the *HSP40* family (Figure 6A-6B).

**Figure 6.**
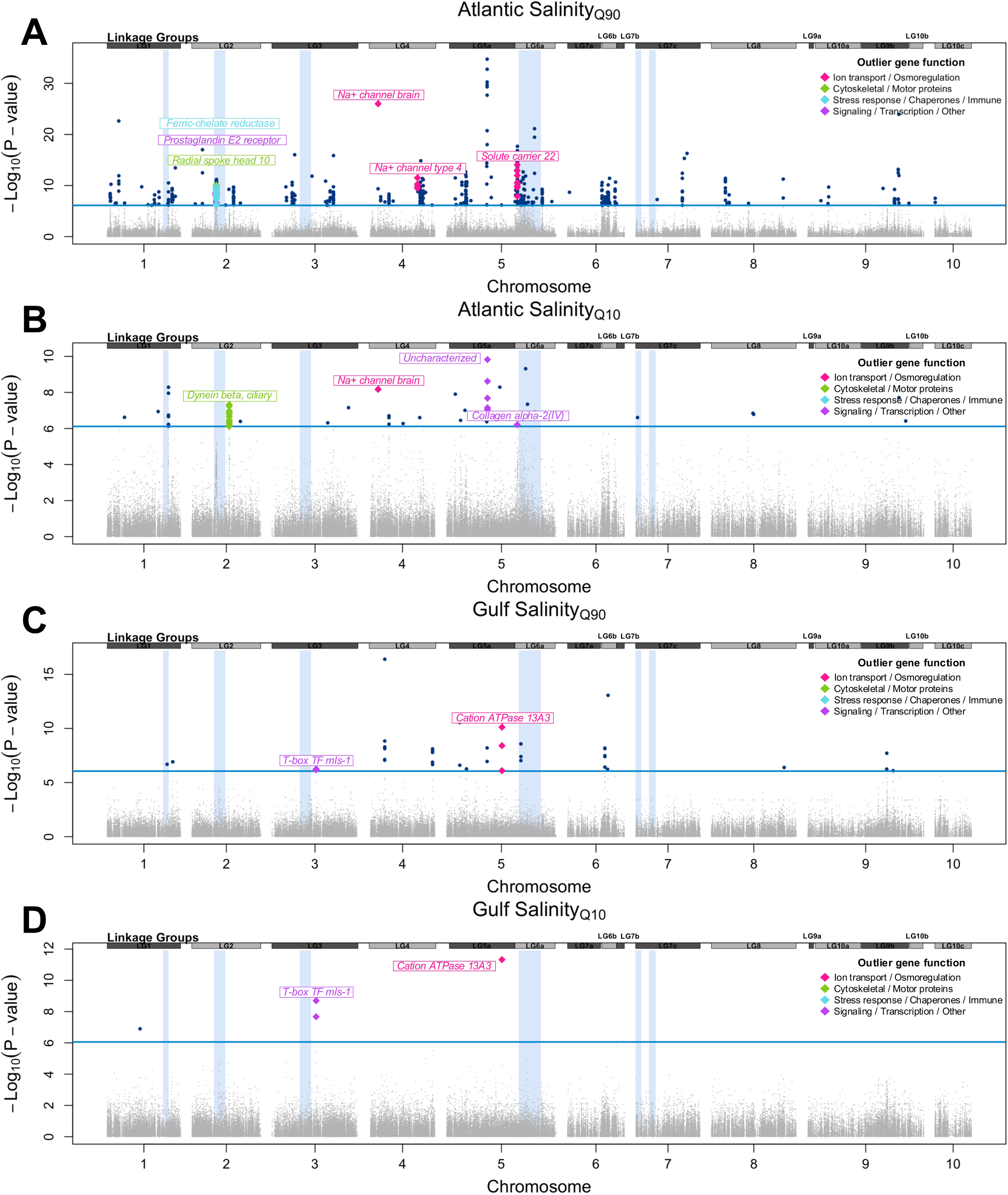
Divergent salinity adaptation in the Gulf and Atlantic populations. A) Manhattan plot of outlier loci (*n* = 814) in association with 90th percentile high salinity (Salinity_Q90_) in the Atlantic. B) Manhattan plot of outlier loci (*n* = 50) in association with 10th percentile low salinity (Salinity_Q10_) outlier loci in the Atlantic. C) Manhattan plot of outlier loci (*n* = 39) in association with 90th percentile high salinity (Salinity_Q90_) outlier loci in the Gulf. D) Manhattan plot of outlier loci (*n* = 5) in association with 10th percentile low salinity (Salinity_Q10_) outlier loci in the Gulf. In all plots (A-D) the linkage groups at the top of the plot correspond to a linkage map published in (Guo et al., 2023) and likely misassemblies in the genome (Puritz et al., 2022, 2024) (Supp. Fig. S5). The blue rectangles in the background correspond to extended LD regions identified by local PCA analysis (Table 2).

Gene ontology (GO) enrichment analysis of the Temperature_Q90_ outlier gene set (*n* = 388) returned significantly enriched functional categories (Supp. Table S4). The pathway with highest fold enrichment was *P-type calcium transporter activity* (GO:0005388). Additional enriched pathways included *9+2 motile cilium* (GO:0097729), *Calcium ion transmembrane transport* (GO:0070588), *Dynein complex* (GO:0030286), and *Iron ion binding* (GO:0005506) (Supp. Table S4).

### Salinity

Genotype-environment association analysis (lfmm2) run separately for the Gulf (220 individuals) and Atlantic (387 individuals) populations identified different sets of loci associated with salinity in each geographic region. In the Atlantic we identified 14 outlier loci corresponding to 167 annotated genes associated with Salinity_Q90_ (Figure 6C) and 50 outlier loci corresponding to 22 annotated genes associated with Salinity_Q10_ (Figure 6D). In contrast, the Gulf had substantially fewer salinity-associated loci, as 39 outlier loci corresponding to 15 annotated genes were associated with 90th quantile high salinity (Salinity_Q90_) (Figure 6E) and 5 outlier loci corresponding to 3 annotated genes were associated with 10th quantile low salinity (Salinity_Q10_) (Figure 6F).

Notably, across all salinity analyses only two loci (of 882 total outlier SNPs = 0.2%) corresponding to 2 (of 191 = 0.1%) genes were shared by both Gulf and Atlantic populations: *Transient receptor potential cation channel subfamily M member 8-like* (LOC111134529: Atlantic Salinity_Q90_ max −log_10_(*P*) = 10.82 and Gulf Salinity_Q90_ max −log_10_(*P*) = 8.57) within the extended LD region on chromosome 5, and an uncharacterized locus also on chromosome 5 (LOC111135484: Atlantic Salinity_Q90_ max −log_10_(*P*) = 34.74, Atlantic Salinity_Q10_ max −log_10_(*P*) = 9.84, and Gulf Salinity_Q90_ max −log_10_(*P*) = 10.99) (Supp. Tables S6-S9). There were no Salinity_Q10_ outliers that were shared between Gulf and Atlantic populations.

Gene ontology enrichment analysis of salinity lfmm2 outlier genes also revealed differences in Gulf and Atlantic-specific significantly enriched (FDR < 0.05) pathways (Supp. Table S3). Atlantic salinity genes were significantly enriched for ion-related pathways, particularly sodium channels, intracellular signaling, and cilium movement. Atlantic high salinity genes were enriched for *Extracellular matrix structural constituent* (GO:0005201), *Intracellular signal transduction* (GO:0035556), *Sodium ion transport* (GO:0006814), *Collagen trimer* (GO:0005581), and *Ion channel activity* (GO:0005216) (Supp. Table S3). Atlantic low salinity genes were enriched for *Cilium movement* (GO:0003341), *Axonemal dynein complex* (GO:0005858), *Cation channel activity* (GO:0005261), *Dynein complex* (GO:0030286), and *Sodium ion transmembrane transport* (GO:0035725) (Supp. Tables S6-S7).

In contrast, Gulf high salinity genes were significantly enriched for cation transporters and transmembrane transport (Supp. Table S3), including *Transmembrane transport* (GO:0055085), *ATPase-coupled cation transmembrane transporter activity* (GO:0019829), *P-type transmembrane transporter activity* (GO:0140358), and *Ion transmembrane transport* (GO:0034220). Gulf low salinity genes were enriched for *ATPase-coupled cation transporter pathway* (GO:0019829) and *Cation transmembrane transport* (GO:0098655) (Supp. Tables S8-S9).

### Spatially heterogeneous selection

Principal-component based outlier detection with pcadapt performed on the global dataset of all individuals (*n* = 745) returned 5524 Bonferroni-corrected outlier SNPs corresponding to 4075 annotated genes (Figure 7A). A strong signal of differentiation was observed on chromosome 7, where two dense clusters of outlier SNPs reached −log_10_(*P*) >150 (*eLD-7a* with 223 outlier SNPs; *eLD-7b* with 129 outlier SNPs, Figure 7A, Table 2). Within *eLD-7a* (Table 2), 55 outlier SNPs were detected in *Negative elongation factor D* (LOC111102475, max −log_10_(*P*) = 220.1), which regulates RNA binding and was also identified in temperature-associated lfmm2 analysis. Within *eLD-7b* (12-18 Mb, LG7c, Table 2), 1 outlier SNP was detected in *Hemagglutinin/amebocyte aggregation factor* (LOC111105403, −log_10_(*P*) = 11.0), a pattern recognition receptor with known immune function (Wang et al., 2025). This locus was also detected as a diagnostic locus with high F_ST_ introgressing in the Chesapeake Bay group. On chromosome 3, *Dynein heavy chain 8* (LOC111123434), responsible for microtubule movement, contained 220 outlier SNPs (max −log_10_(*P*) = 20.5). On chromosome 8, 41 outlier SNPs were detected in *Voltage-dependent calcium channel subunit alpha type D* (LOC111110350, max −log_10_(*P*) = 100.03), which enables high-voltage calcium ion transport across the plasma membrane.

**Figure 7.**
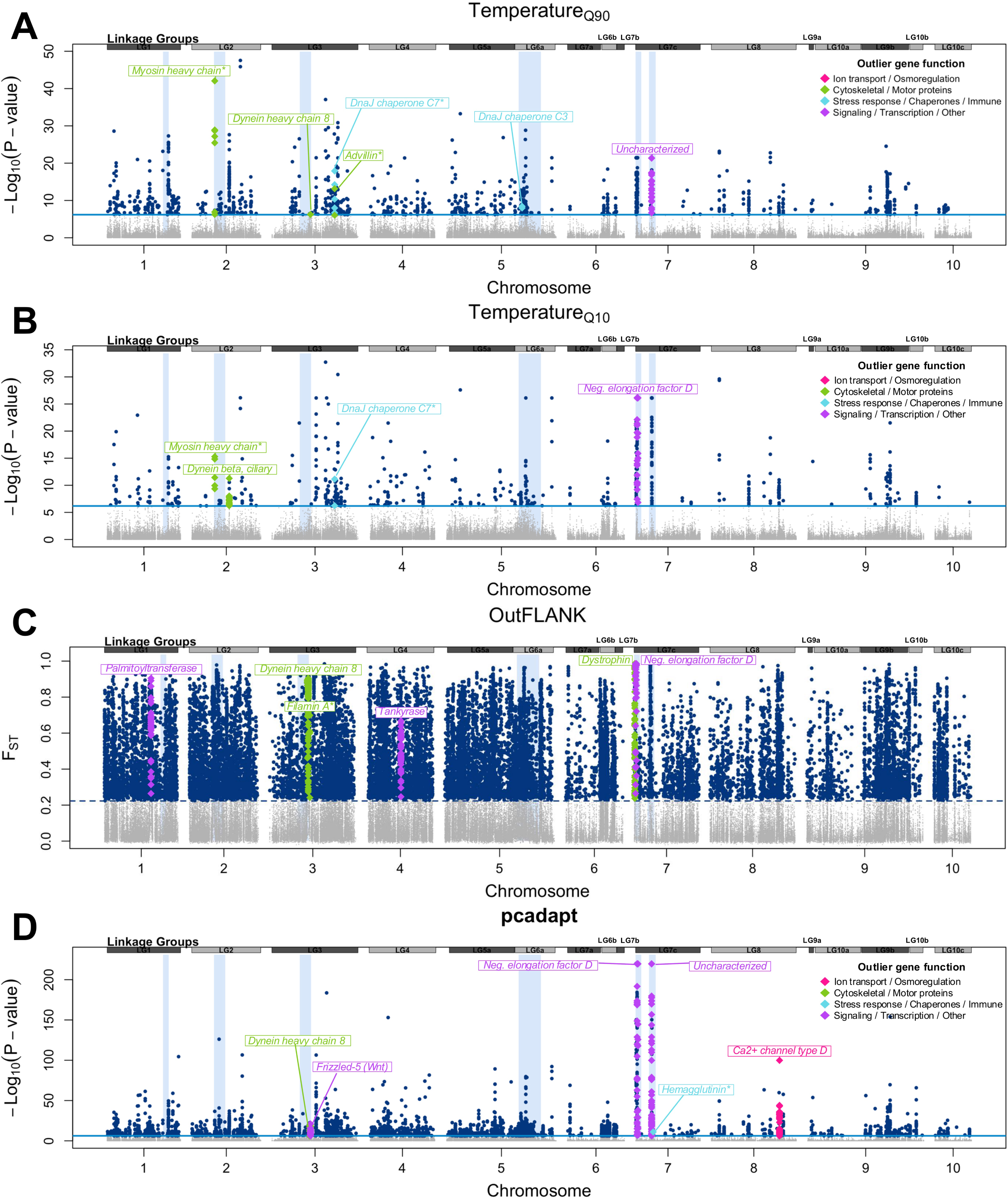
Signatures of spatially heterogeneous selection and temperature adaptation across the range. A) Manhattan plot of outlier loci (*n* = 1318) in association with global 90th percentile high temperature (Temperature_Q90_). B) Manhattan plot of outlier loci (*n* = 734) in association with global 10th percentile low temperature (Temperature_Q10_). C) Manhattan plot of outlier loci detected by OutFLANK (*n* = 23,410). Genes colored in green have cytoskeletal or motor protein function, genes colored in purple are involved in signaling, transcription, or other functions. D) Manhattan plot of outlier loci detected by pcadapt (*n* = 5,524). In all plots (A-D) the linkage groups at the top of the plot correspond to a linkage map published in (Guo et al., 2023) and likely misassemblies in the genome (Puritz et al., 2022, 2024) (Supp. Fig. S5). The blue rectangles in the background correspond to extended LD regions identified by local PCA analysis (Table 2).

**Figure 8.**
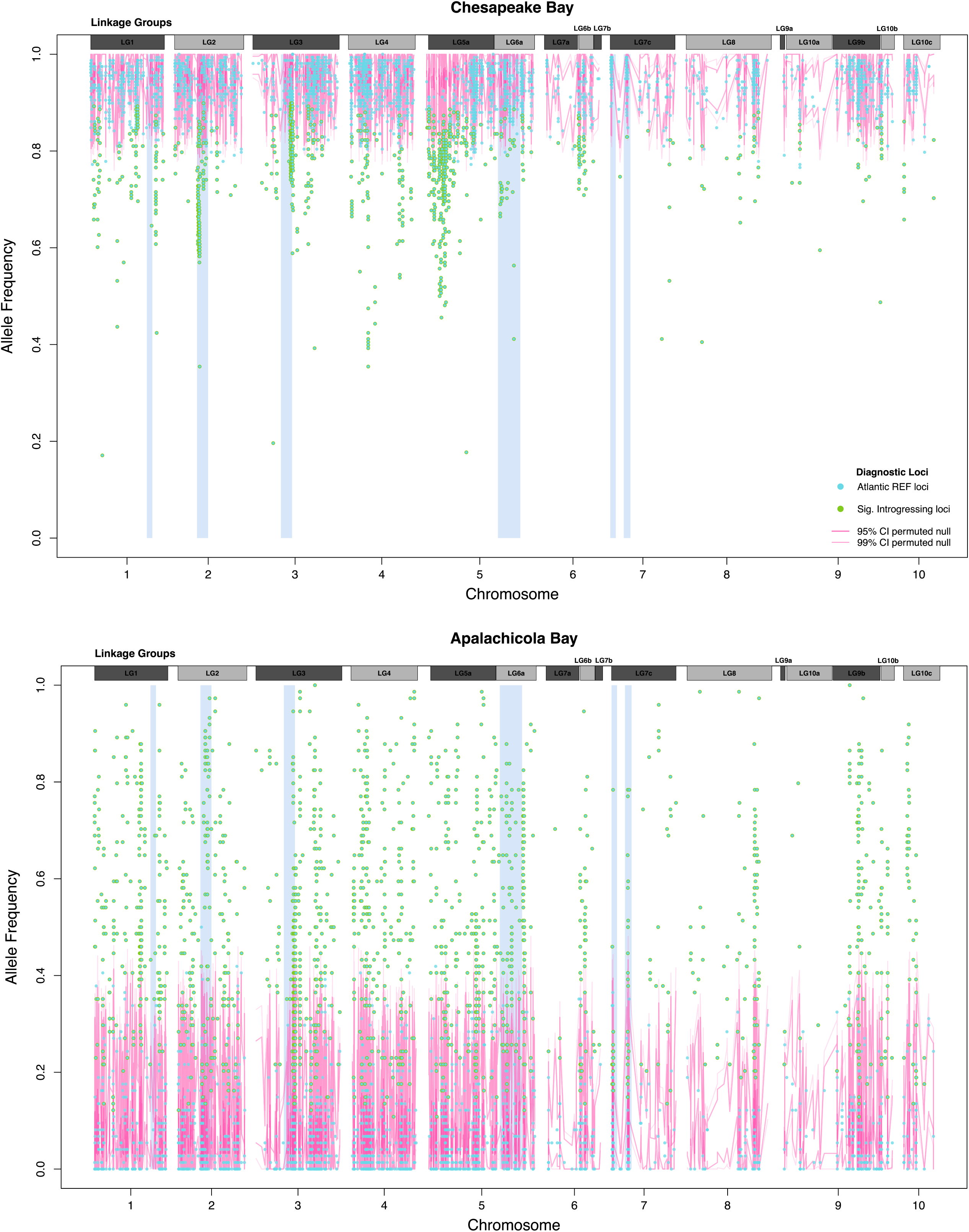
Introgression at diagnostic loci across the genome. Allele frequency of Atlantic (REF) diagnostic loci (*n* = 5,542) across the genome in (A) the admixed Chesapeake Bay sites and (B) the admixed Apalachicola Bay sites. Loci in blue were not detected as significantly introgressing by permutation testing. Loci outlined in green were detected as significantly introgressing by permutation testing (FDR < 0.01) with a change in allele frequency more than 0.1 (Δp < - 0.1 in Chesapeake Bay, Δp > 0.1 in Apalachicola Bay). Permuted null confidence intervals (95% and 99%) are in pink. Linkage groups at the top of the plot correspond to a linkage map published in (Guo et al., 2023) and likely misassemblies in the genome (Puritz et al., 2022, 2024) (Supp. Fig. S5). The blue rectangles in the background correspond to extended LD regions identified by local PCA analysis (Table 2).

Gene ontology (GO) enrichment analysis of the pcadapt outlier gene set returned 172 significantly enriched (FDR < 0.05) functional categories (Supp. Table S10). The two pathways with highest fold enrichment were *L-absorbic acid transmembrane transporter activity* (GO:0015229, *n* = 17 genes) and *L-absorbic acid transmembrane transport* (GO:0015882; *n* = 17 genes). Additional enriched pathways included *Chloride transmembrane transport* (GO:1902476), *Minus-end-directed microtubule motor activity* (GO:0008569), and *Dynein complex* (GO:0030286) (Supp. Table S10).

Outlier detection with OutFLANK performed on the global dataset of all individuals (*n* = 745) returned 23,410 outlier SNPs corresponding to 6194 annotated genes (Figure 7B). Similarly to pcadapt results, 245 outliers (max −log_10_(*P*) = 3.6) were located within *Dynein heavy chain 8* (LOC111123434) within the extended LD region on chromosome 3 (eLD-3 in Table 2). Within eLD-3, *Filamin A-interacting protein 1-like* (LOC111123901) contained 5 outlier loci (max −log_10_(*P*) = 2.73). On chromosome 1, *Palmitoyltransferase ZDHHC17-like, transcript variant X1* (LOC111124769) had 58 outlier loci (max −log_10_(*P*) = 3.61), and on chromosome 4, *Tankyrase-like protein* (LOC111129603) had 68 detected outliers (max −log_10_(*P*) = 2.62). Within extended LD region eLD-7a (Table 2), genes *Negative elongation factor D-like transcript variant X2* (n = 59 outliers) and *Dystropin-like* (*n* = 76 outliers) each had a high number of significant outliers (max −log_10_(*P*) = 2.62).

Gene ontology enrichment analysis of the OutFLANK outlier gene set (*n* = 6194) returned 319 significantly enriched (FDR <0.05) functional categories (Supp. Table S3). The pathway with highest fold enrichment was *Heme binding* (GO:0020037; *n* = 75 genes). Other highly enriched pathways included *DNA-binding transcription factor activity* (GO:0003700; *n* = 295 genes) and *Transmembrane transport* (GO:0055085; *n* = 730 genes) (Supp. Table S11).

### Functional Enrichment of Consensus Genes

To identify genes with the most consistent evidence of selection, we ranked all annotated genes by the number of independent analyses in which they contained at least one outlier SNP (Supp. Table S12). There were a maximum of 10 relevant analyses from which we extracted outlier SNP information (pcadapt, OutFLANK, two temperature lfmm2 tests, four salinity lfmm2 tests, and two introgression tests). Eight genes were detected in 7 out of 10 analyses, and an additional 67 genes were detected in at least 6 analyses (Supp. Table S3).

The highest ranked consensus gene was *Negative elongation factor D-like, transcript variant X2* (LOC111102475, chromosome 7, 56 SNPs, 7 analyses), a transcriptional regulator involved in RNA polymerase II pausing, which is located within extended LG region *eLD-7a* on chromosome 7 (Table 2). Three other genes that were detected in at least 7 analyses were also located within *eLD-7a*: *Ashwin-like transcript variant X5* (LOC111104055, chromosome 7, 21 SNPs), an uncharacterized locus (LOC111104062, chromosome 7, 13 SNPs), and *Xanthine dehydrogenase-like* (LOC111136587, chromosome 7, 12 SNPs). Other prominent consensus genes included *Probable G-protein coupled receptor B0563.6* (LOC111120197, 13 SNPs) and *Ferric-chelate reductase 1-like transcript variant X2* (LOC111121220, 28 SNPs). *Dynein heavy chain 8*, located within extended LG region *eLD-3* (Table 2) was detected in 6 analyses with 225 outlier SNPs. *Voltage-dependent calcium channel type D* (LOC111110350, chromosome 8, 42 SNPs, 6 analyses), *Dystrophin* (LOC111103558, chromosome 7, 41 SNPs, 6 analyses), and *Frizzled-5* (LOC111122602, chromosome 3, 49 SNPs, 6 analyses) were among other top consensus genes detected in at least 6 analyses.

GO enrichment analysis of the top 100 consensus genes revealed functional categories dominated by cytoskeletal motor activity and dynein-related pathways, including *Dynein complex* (GO:0030286), *Minus-end-directed microtubule motor activity* (GO:0008569), *Microtubule-based movement* (GO:0007018), and *Actin filament binding* (GO:0051015). *Calcium ion transmembrane transport* (GO:0070588), *Lipid metabolism* (GO:0006629), *G protein-coupled receptor signaling* (GO:0007186), and *Translational elongation* (GO:0006414) pathways were also significantly enriched.

## Discussion

This study presented a range-wide population genomic survey of wild eastern oysters. We identified two primary ancestral groups (Gulf and Atlantic), with finer-scale substructure in each region. We also discovered unexpected patterns of ancestry in the Chesapeake Bay/Pamlico Sound and Apalachicola Bay, which we discuss potential causes of below. We identified regions of extended linkage disequilibrium across four chromosomes, some of which harbored strong outlier signals in genome scans and may play an important role in adaptation. Genotype-environment association analyses revealed largely independent pathways of salinity adaptation between Gulf and Atlantic groups and also multiple temperature-associated genes previously implicated in disease resistance. Collectively, these findings demonstrate that the genomic landscape of the eastern oyster is shaped by an interplay of environmental selection, structural variation, introgression, and direct and indirect human effects of management.

### Population Structure and Divergence

We identified two primary ancestral groups, consistent with previous studies that identified differentiation between eastern oysters in the Gulf from their allopatric conspecifics on the Atlantic coast using mtDNA markers (Reeb & Avise, 1990), nuclear DNA RFLPs (Hoover & Gaffney, 2005; Karl & Avise, 1992), and more recent SNP-based assessments (Puritz et al., 2022; Thongda et al., 2018). The scale of differentiation between Gulf and Atlantic populations, evidenced by global maximum pairwise F_ST_ = 0.0647 between Calcasieu Lake, Louisiana (site 6) and Prince Edward Island, Canada (site 40), suggests long-term isolation maintained by oceanographic features with the Florida peninsula south of Cape Canaveral as a barrier to gene flow (Hare & Avise, 1996; Murray & Hare, 2006). Within these two primary groups, we also detected patterns of regional substructure, particularly among Gulf populations.

Within the Gulf, we observed different genetic partitioning than previously detected with other marker types; the earliest work to differentiate populations of eastern oysters found no significant allozyme differentiation within Gulf populations (Buroker, 1983). A more recent study determined oyster populations in the panhandle of Florida to be part of the Gulf metapopulation using mtDNA and nuclear SNPs (Varney et al., 2009). In our study, clear partitioning was detected between Apalachicola Bay and other Gulf sites in Alabama, Louisiana, and Texas, which collectively formed a single genetic cluster, a notable deviation from previous studies. Prior work identified two genetically differentiated populations between Laguna Madre, TX and populations to the north in Texas (Anderson et al., 2014), and this genetic divergence may explain the genotyping failure of Laguna Madre individuals in this study.

Within the Atlantic we observed complex patterns of substructure. Sites south of Pamlico Sound (North Carolina) including Georgia, South Carolina, North Carolina, and the Atlantic coast of Florida formed a single genetic cluster. The mid-Atlantic region, spanning from North Carolina’s Pamlico Sound to Long Island sound (NY), showed patterns of admixture consistent with isolation-by-distance. We found the Chesapeake Bay sites to be genetically similar, which is likely due to a combination of high gene flow and extensive oyster restoration with non-local genotypes (Rose et al., 2006). We discovered regional substructure north of Cape Cod between Southern New England, New Hampshire, Maine, and Canada. We did not detect substructure within Canadian populations, although we did not sample as densely as previous work which described several genetic clusters using RADseq in Canadian populations (Bernatchez et al., 2019). Our findings expand upon previously detected subtle structure along the Atlantic coast (Puritz et al., 2022; Rose et al., 2006) by providing increased resolution of differences among Atlantic populations.

Notably, F_IS_ values were negative across all sampling locations (Supp. Table S1). Heterozygote excess is common in wild populations of highly fecund marine bivalves and may reflect a combination of high genetic load and sweepstakes reproductive success characteristic of broadcast spawning marine invertebrates (Hedgecock & Pudovkin, 2011; Plough, 2016). Negative F_IS_ values have also been reported in wild and restored eastern oyster populations in Great Bay, NH from a prior study which used low-coverage whole genome sequencing (Strickland & Brown, 2024); our study reports a similar result from that same estuary with the SNP array (Great Bay, NH F_IS_ = -0.2434, Supp. Table S1).

### Human impacts and unexpected geographic patterns of ancestry

We discovered unexpected patterns of Gulf ancestry in the mid-Atlantic (Chesapeake Bay, VA and Pamlico Sound, NC) and Atlantic ancestry in the eastern Gulf (Apalachicola Bay, FL). These patterns resembled introgression with multiple generations of backcrossing, but in this two-cluster scenario similar patterns could also result from genetic drift (i.e., when alleles at high frequency in an ancestral cluster drift, the only direction they can drift without being lost is toward the other ancestral cluster). While we cannot conclusively determine the ultimate causes of these patterns, both the greater Chesapeake area and Apalachicola Bay have an intense history of human impacts that may have strongly influenced these unexpected geographic patterns of ancestry.

Apalachicola showed admixed Atlantic and Gulf ancestry and was genetically distinct from other Gulf sites. Gene flow from Atlantic populations around the Florida peninsula could introduce Atlantic alleles, however, previous studies using mtDNA and microsatellites did not detect Apalachicola Bay as genetically distinct from other Gulf populations (Varney et al., 2009), and the Florida peninsula has historically been considered a strong biogeographic barrier to gene flow (Murray & Hare, 2006). The introgression and genetic divergence we detected may reflect genetic drift following demographic decline. Apalachicola Bay experienced a severe population collapse over the past decade, driven by a combination of drought-induced salinity changes, fishing pressure, periodic droughts, and disease (Hintenlang et al., 2023; Lawrance et al., 2017; Pine et al., 2015). Starting as early as 2007, reduced freshwater flow from the Apalachicola-Chattahoochee-Flint river system due to changes to upstream water allocation increased salinity of the bay (Oczkowski et al., 2011; Petes et al., 2012). In turn, this intensified *P. marinus* infection in Apalachicola Bay and oyster predation (Pusack et al., 2019), causing significant oyster declines to the point of fishery collapse in 2012 (Hintenlang et al., 2023; Pine et al., 2015). The resulting population bottleneck may have shifted allele frequencies through drift, producing ancestry estimates that are divergent from the larger Gulf population, we observed genome-wide shifts in frequency consistent with this expectation. Future work that includes more samples around the Florida peninsula, as well as historical samples from Apalachicola, could help elucidate the roles of gene flow and genetic drift in driving genetic diversity in this region.

The unexpected pattern of Gulf ancestry in the Chesapeake Bay was interesting in its geographic isolation, without similar patterns to the south. This suggests that if the pattern is due to introgression, it is due to artificial gene flow (*e.g.,* human movement of oysters from the Gulf to the Chesapeake Bay) rather than natural dispersal. Indeed, the Chesapeake Bay has a documented history of importing Gulf oysters after the MSX epidemic in the early 1960s, which supplied oyster shucking in Virginia for multiple subsequent decades (Schulte, 2017). Within the past three decades, Gulf oysters from Louisiana have also been transported to the Chesapeake region to develop and breed founder populations. In the Chesapeake, we observed some regions with a concentration of introgressing loci, particularly on chromosome 5 and within structural variants on chromosomes 2 and 3, although there were also additional signals spread broadly across the genome. The consequences of this genetic mixing is unclear. On one hand, translocation and admixture between highly differentiated populations could reduce inheritance of locally adapted alleles.

This process, known as ‘genetic swamping,’ has been documented as a risk to aquaculture (Roberts et al., 2010; Waples et al., 2012). Similarly, outbreeding depression can reduce population fitness by disrupting inheritance of co-adaptive gene complexes (Aitken & Whitlock, 2013; Waples et al., 2012). For instance, introgression from farmed to wild salmon has been shown to alter genetic diversity and reduce fitness (San Román et al., 2025), and disrupt locally adapted gene complexes (Glover et al., 2017). On the other hand, introduction of southern alleles into a northern population is consistent with what an assisted gene flow program (Aitken & Whitlock, 2013) would propose to increase thermal tolerance of the northern population. A promising area of future research is determining the fitness consequences and adaptive potential of introgressing Gulf alleles into the Chesapeake.

### Putative Structural Variants

Structural variants are increasingly recognized as drivers of adaptive variation because they suppress recombination and preserve adaptive gene complexes even under high gene flow (Schaal et al., 2022; Tigano & Friesen, 2016; Wellenreuther & Bernatchez, 2018). We detected six putative inversion regions across four chromosomes identified through local PCA analysis. Several highly ranked consensus genes detected in our study were located within structural variant regions. As discussed in more depth in the next section, clusters of outlier loci within structural variants may be an important implication for wild adaptation and aquaculture breeding.

### Environmental Adaptation

#### Salinity

In the eastern oyster, genotype-by-environment interactions have been documented by multiple studies, including population-specific transcriptomic responses to low salinity (Johnson & Kelly, 2020; Sirovy et al., 2023), heritable genetic variation for low salinity survival (McCarty et al., 2022), and polygenic salinity-associated loci distributed across the genome (Bernatchez et al., 2019). Our results expand upon these prior findings with a range-wide dataset to reveal that Gulf and Atlantic ancestral populations have differentially adapted to salinity stress, although we note reduced power from a smaller sampling size in the Gulf. Adaptive divergence in osmoregulatory gene expression has previously been reported in other *Crassostrea* species under different salinity gradients (Zhang et al., 2023).

Of the salinity-associated outlier SNPs we identified across genotype-environment association analyses, few were shared between Gulf and Atlantic groups. Although both groups showed evidence of enrichment for transmembrane transport and ion-related GO pathways, this was achieved through different underlying genes. The lack of overlap at the SNP, gene, and functional pathway levels is consistent with divergent adaptation (Popovic & Riginos, 2020; Zhang et al., 2023), as the two lineages have independently evolved different osmoregulatory mechanisms.

Genes implicated in low salinity tolerance were also detected in other analyses. Our study found several *Dynein heavy chain* genes and a significantly enriched *Dynein complex* pathway across multiple analyses. *Dynein heavy chains* are molecular motor proteins that are involved with microtubule-based movement, cilium function, ATP binding, cytoplasm and cytoskeleton. For instance, *Dynein heavy chain 8*, located in extended LD region *e-LD3* contained the largest signal from spatially heterogeneous selection analyses, and was also detected as a temperature-associated outlier in both high and low temperature genotype-environment association analyses. Further, all outliers in *Dynein heavy chain 8* were detected as diagnostic loci, and many showed significant introgression into the Chesapeake Bay and Apalachicola Bay. We also found *Dynein beta chain, ciliary-like* to contain the most outlier SNPs from Atlantic low salinity analysis. In *Palmitoyltransferase ZDHHC17*, the majority of SNPs were OutFLANK F_ST_ outliers diagnostic loci with significant introgression. One previous study found *Dynein heavy chain* and *Palmitoyltransferase ZDHHC17* to be hypermethylated in oysters from a low salinity Gulf site (Johnson & Kelly, 2020). *Palmitoyltransferase ZDHHC17* catalyzes fatty acids to proteins, which is critical for protein membranes and signaling. While (Johnson & Kelly, 2020) found that genetic differences among Gulf populations alone could not explain hypermethylation patterns at these genes, our range-wide sampling in both the Gulf and Atlantic has revealed that *Dynein* and *Palmitoyltransferase* genes show clear genetic differentiation between the two ancestral groups. These genes are clearly associated with environmental gradients and are actively introgressing, contributing to adaptation across the range of eastern oysters.

#### Temperature and Disease

Our analyses identified genes involved in environmental adaptation that have also been previously implicated in resistance to the parasite *Perkinsus marinus (Wang et al., 2025)*, the causative agent of Dermo disease. Parasite proliferation is strongly temperature-dependent; *P. marinus* replication accelerates at high temperatures, infection intensity peaks during or shortly after warm summer months, and northward range expansion of Dermo has been linked to increasing sea surface temperatures (Cook et al., 1998; La Peyre et al., 2008). High temperatures simultaneously suppress host lysozyme activity and disrupt energy metabolism, compounding the effects of parasitic infection and driving mortality (F. L. E. Chu & La Peyre, 1993; Encomio & Chu, 2007).

Of the overlapping genes between our range-wide analyses and known Dermo resistance genes (Wang et al., 2025), we identified three genes in association with high temperature: *Dnaj homolog subfamily C member 7*, a heat shock protein (HSP) in the HSP40 family that co-chaperones HSP70 to refold damaged proteins under thermal stress; *Myosin heavy chain, non-muscle like*, associated with HSP90 chaperone activity and cytoskeletal integrity; and *Advillin*, an actin-binding protein involved in cytoskeletal dynamics and hemocyte phagocytosis. Genes related to both Dermo resistance and high-temperature adaptation are biologically significant as they may pose an adaptive response to dual stressors. We detected an additional two genes previously implicated in Dermo resistance (Wang et al., 2025) under spatially heterogeneous selection analyses: *Hemagglutinin/amebocyte aggregation factor*, involved in pathogen opsonization, and *Filamin A-interacting protein 1*. Both of these genes contained a diagnostic locus which we detected as significantly introgressing into the Chesapeake Bay group. These findings have practical implications for aquaculture breeding programs, as genomic selection for Dermo resistance has been noted as having slow progress (Guo, 2021; Powell et al., 2011; Wang et al., 2025). Overlap between genes which are adaptive to temperature stress and involved in Dermo resistance may be subject to balancing selection across heterogeneous environments (Yu & Guo, 2006). As climate change intensifies thermal stress and disease pressure (Cook et al., 1998; Ford, 1996), understanding overlap of adaptive genes is crucial for predicting adaptive capacity in wild populations and for designing multi-trait genomic selection programs for aquaculture.

Functional enrichment analysis of consensus genes identified by both spatially heterogeneous selection and environmental selection further support the role of immune-related pathways in adaptation. We detected enrichment for *Lipase activity* and *Triglyceride lipase activity*, involved in immune signaling and mobilization of lipid mediators during pathogen challenges (F.-L. E. Chu et al., 2002; Pernet et al., 2007). We also found highly enriched pathways for *L-absorbic acid transmembrane transporter activity* and *transport*, important components of antioxidant defense which mitigate oxidative damage in bivalves exposed to bacterial challenges including *Vibrio* infections (Genard et al., 2013; Luo et al., 2021). These findings suggest selection on immune function may be an important driver of adaptation in eastern oysters across the seascape, and future studies should investigate finer-scale associations between regional genetics and disease gradients, as well as the relationship between underlying genotype and parasite load.

We acknowledge that GO analyses are subject to limitations and biases, including incomplete SNP annotation and a bias towards well-characterized gene families (Tiffin & Ross-Ibarra, 2014). However, our mapping of additional annotations onto the existing eastern oyster genome substantially improved functional resolution and we suggest future genetic analyses re-annotate to improve functional resolution of enriched targets. Hence, despite these limitations, our findings provide valuable insights into the genomic architecture of adaptation in eastern oysters and highlight the complex interplay between environmental selection and introgression in shaping variation across the seascape.

### Conclusions

Globally expanding capacity in marine aquaculture presents new opportunities to establish food security and meet demand for sustainably-produced protein. In the United States, the seafood trade deficit stands at nearly $20 billion, and advancing selective breeding of native aquaculture species is a priority for closing this gap (Andersen et al., 2025; Houston et al., 2020; Naylor et al., 2021). Our range-wide seascape genomic study of wild eastern oyster populations provides actionable insights for integrating wild population diversity into aquaculture for improvement. Climate change is intensifying thermal stress and disease pressure, shifting the environmental landscape for wild and cultured populations alike and dictating priorities for aquaculture improvement. Our findings provide valuable insights into the genomic architecture of adaptation in eastern oysters, highlighting the complex interplay between environmental selection and introgression in shaping variation across the seascape and demonstrating how integrating seascape genomics into aquaculture breeding can advance both conservation, management and aquaculture production goals.

## Data Accessibility

All code and data are available for peer review at the GitHub site https://github.com/ModelValidationProgram/OysterSeascape. Upon final acceptance, data will be archived on BCO-DMO in the project “CAREER: Evaluation of machine learning algorithms for understanding and predicting adaptation to multivariate environments with a Model Validation Program (MVP)” available at https://www.bco-dmo.org/project/876610.

