## Supplement for "Range-wide genetic population structure and environmental adaptation in the eastern oyster (*Crassostrea virginica*) provides insight for aquaculture"

Supplemental Tables 2

Supp. Table S1: Ancestry and genetic diversity for each sampling location 2

Supp. Table S2: Environmental conditions at each sampling location 3

Supp. Table S3: Mutation table with all SNPs and significance values from all analyses 4

Supp. Table S4: lfmm2 global high temperature enrichment analysis 4

Supp. Table S5: lfmm2 global low temperature enrichment analysis 4

Supp. Table S6: lfmm2 Atlantic high salinity enrichment analysis 4

Supp. Table S7: lfmm2 Atlantic low salinity enrichment analysis 4

Supp. Table S8: lfmm2 Gulf high salinity enrichment analysis 4

Supp. Table S9: lfmm2 Gulf low salinity enrichment analysis 4

Supp. Table S10: pcadapt gene ontology enrichment analysis 4

Supp. Table S11: OutFLANK gene ontology enrichment analysis 4

Supp. Table S12: Top 100 consensus genes 5

Supplemental Figures 5

Fig. S1. Global snmf cross-entropy plot 5

Fig. S2. Gulf snmf cross-entropy plot 6

Fig. S3. Atlantic snmf cross-entropy plot 7

Fig. S4. Local PCA MDS1 window plot for all chromosomes 8

Fig. S5. Synteny plot with chromosome-linkage group misalignments 9

### *Supplemental Tables*

#### *Supp. Table S1: Ancestry and genetic diversity for each sampling location*

Summary of sampling sites, genetic ancestry, and diversity statistics for 40 sampling locations of eastern oysters spanning the Gulf (southern Texas) to Atlantic Canada. Sites are numbered geographically from south to north. Genetic cluster assignments are based on ancestry analysis (Figure 1, main text), with proportions (*Q1 Gulf*, *Q2 Atlantic*) representing the mean proportion of ancestral cluster assignment from all sampled individuals at each site. Laguna Madre, TX (Site 1) was excluded from analysis due to low genotyping success rate.

| Site | Region | State | Lat/Long | Genetic Cluster | Q1 Gulf | Q2 Atlantic | Ho | He | $F_{IS}$ |
|---|---|---|---|---|---|---|---|---|---|
| 1 | Laguna Madre | TX | 26.5597, -97.3591 | - | - | - | - | - | - |
| 2 | Copano Bay | TX | 28.0960, -97.1740 | Gulf | 0.98720961 | 0.01279042 | 0.4157 | 0.297 | -0.2841 |
| 3 | Tres Palacios Bay | TX | 28.6936, -96.2233 | Gulf | 0.98457995 | 0.01542006 | 0.4163 | 0.2975 | -0.2826 |
| 4 | Galveston Bay | TX | 29.5475, -94.9036 | Gulf | 0.93861032 | 0.06138965 | 0.4145 | 0.2985 | -0.2696 |
| 5 | Sabine Lake | LA | 29.785, -93.9180 | Gulf | 0.99378526 | 0.00621482 | 0.4166 | 0.2973 | -0.2848 |
| 6 | Calcasieu Lake | LA | 29.8449, -91.3179 | Gulf | 0.9950603 | 0.00493974 | 0.4156 | 0.2965 | -0.2855 |
| 7 | Cote Blanche Bay | LA | 29.4196, -91.7071 | Gulf | 0.99366732 | 0.00633264 | 0.4164 | 0.2974 | -0.2839 |
| 8 | Sister Lake | LA | 29.2399, -90.9113 | Gulf | 0.99549863 | 0.00450135 | 0.4162 | 0.2974 | -0.2835 |
| 9 | Barataria Bay | LA | 29.4236, -90.0101 | Gulf | 0.99392463 | 0.00607542 | 0.4166 | 0.2975 | -0.2846 |
| 10 | Lake Fortuna | LA | 29.6437, -89.4976 | Gulf | 0.99320183 | 0.00679823 | 0.4159 | 0.2968 | -0.2851 |
| 11 | Grand Bay | AL | 30.3727, -88.3183 | Gulf | 0.99129429 | 0.00870565 | 0.4155 | 0.2967 | -0.285 |
| 12 | Saint Vincent Sound | FL | 29.6891, -85.2207 | Apalachicola | 0.8826525 | 0.1173475 | 0.4125 | 0.2962 | -0.277 |
| 13 | East Cove | FL | 29.6956, -84.7893 | Apalachicola | 0.87979684 | 0.12020316 | 0.413 | 0.2963 | -0.2768 |
| 14 | Timucuan Preserve | FL | 30.4400, -81.4363 | Atlantic | 0.0325361 | 0.96746395 | 0.3894 | 0.2874 | -0.2349 |
| 15 | Sapelo Island | GA | 31.4177, -81.2957 | Atlantic | 0.03349668 | 0.96650339 | 0.389 | 0.2881 | -0.232 |
| 16 | Doboy Sound | GA | 31.4535, -81.3631 | Atlantic | 0.03278193 | 0.9672181 | 0.3887 | 0.2878 | -0.2304 |
| 17 | Skidaway River | GA | 31.9899, -81.0217 | Atlantic | 0.0297357 | 0.97026439 | 0.3881 | 0.2871 | -0.2327 |
| 18 | Bull River | GA | 32.0172, -80.9220 | Atlantic | 0.02432647 | 0.9756735 | 0.3883 | 0.2871 | -0.2321 |
| 19 | Charleston Harbor | SC | 32.7523, -79.8976 | Atlantic | 0.02008866 | 0.97991137 | 0.388 | 0.2868 | -0.2334 |
| 20 | Murrells Inlet | SC | 33.5235, -79.0618 | Atlantic | 0.01998049 | 0.98001942 | 0.3877 | 0.2867 | -0.2328 |
| 21 | Cape Fear River | NC | 33.9586, -77.9418 | Atlantic | 0.0153002 | 0.98469975 | 0.3872 | 0.2865 | -0.2315 |
| 22 | Masonboro Island | NC | 34.1402, -77.8636 | Atlantic | 0.01533548 | 0.98466463 | 0.3865 | 0.2858 | -0.2321 |
| 23 | Pamlico Sound | NC | 35.7938, -75.5497 | Chesapeake | 0.08311299 | 0.91688711 | 0.39 | 0.2909 | -0.2245 |
| 24 | Wreck Shoal | VA | 37.0638, -76.5627 | Chesapeake | 0.0816597 | 0.9183404 | 0.3883 | 0.2895 | -0.224 |
| 25 | Deep Water Shoal | VA | 37.1488, -76.6355 | Chesapeake | 0.1204041 | 0.87959602 | 0.3905 | 0.2918 | -0.2217 |
| 26 | Eastern Shore | VA | 37.6166, -75.6639 | Atlantic | 0.01503591 | 0.98496416 | 0.3854 | 0.2874 | -0.2253 |
| 27 | Potomac River | MD | 38.3369, -76.9766 | Chesapeake | 0.07777167 | 0.92222842 | 0.3885 | 0.2895 | -0.2248 |
| 28 | Cape Shore | NJ | 39.0735, -74.9130 | Atlantic | 0.02250597 | 0.9774941 | 0.3868 | 0.288 | -0.2262 |
| 29 | Hope Creek | NJ | 39.4347, -75.5169 | Atlantic | 0.02548016 | 0.97451983 | 0.387 | 0.288 | -0.227 |
| 30 | Ash Creek | CT | 41.1464, -73.2357 | Atlantic | 0.01944541 | 0.98055458 | 0.3811 | 0.2849 | -0.2225 |
| 31 | Long Island Sound | CT | 41.2719, -72.5861 | Atlantic | 0.01949912 | 0.98050084 | 0.3818 | 0.2857 | -0.2219 |
| 32 | Narrow River | RI | 41.5050, -71.4530 | Atlantic | 0.03498429 | 0.96501575 | 0.385 | 0.2853 | -0.2306 |
| 33 | Narragansett Bay | RI | 41.5589, -71.4378 | Atlantic | 0.0169536 | 0.98304645 | 0.3852 | 0.287 | -0.2254 |
| 34 | Plum Island Sound | MA | 42.7517, -70.8365 | Atlantic | 0.01408212 | 0.98591785 | 0.3859 | 0.2855 | -0.2329 |
| 35 | Great Bay | NH | 43.0537, -70.9115 | Atlantic | 0.0217899 | 0.97821005 | 0.3852 | 0.2823 | -0.2434 |
| 36 | Orrs Cove | ME | 43.8313, -69.9153 | Atlantic | 0.02111654 | 0.97888345 | 0.3792 | 0.2806 | -0.2342 |
| 37 | Damariscotta River | ME | 44.0245, -69.5320 | Atlantic | 0.01128816 | 0.98871185 | 0.3744 | 0.2783 | -0.2313 |
| 38 | Salt Bay | CN | 43.8124, -65.9105 | Atlantic | 0.00221191 | 0.99778816 | 0.3832 | 0.2799 | -0.2468 |
| 39 | Eel Lake | CN | 43.8274, - 65.9007 | Atlantic | 0.00474878 | 0.99525118 | 0.3838 | 0.2805 | -0.2481 |
| 40 | Prince Edward Island | CN | 46.1459, -62.8874 | Atlantic | 0.00215467 | 0.9978453 | 0.3833 | 0.2797 | -0.2472 |

*Supp. Table S2: Environmental conditions at each sampling location*

| Site | Region | State | Lat of envi. data source | Long of envi. data source | SalinityQ10 | SalinityQ90 | TemperatureQ10 | TemperatureQ90 | Years | Data Source |
|---|---|---|---|---|---|---|---|---|---|---|
| 1 | Laguna Madre | TX | - | - | - | - | - | - | - | - |
| 2 | Copano Bay | TX | 28.0841 | -97.2009 | 5.9 | 36.7 | 13.8 | 30.5 | 16 | NERR Centralized Data. Mission Aransas - Copano Bay West MARCWWQ |
| 3 | Tres Palacios Bay | TX | 28.6865 | -96.2398 | 7.0 | 27.0 | 13.2 | 30.9 | 3 | USGS Water Data Tres Palacios Bay - 08162675 |
| 4 | Galveston Bay | TX | 29.6703 | -94.8536 | 1.0 | 17.0 | 12.2 | 30.0 | 3 | USGS Water Data Galveston Bay - 08067303 |
| 5 | Sabine Lake | LA | 29.7580 | -93.8900 | 2.6 | 23.8 | 12.6 | 30.6 | 32 | Water Data For Texas - Lower Sabine |
| 6 | Calcasieu Lake | LA | 29.8157 | -93.3490 | 8.7 | 25.0 | 13.4 | 30.7 | 16 | USGS Water Data Calcasieu River - 08017118 |
| 7 | Cote Blanche Bay | LA | 29.7132 | -91.8803 | 0.4 | 8.3 | 11.7 | 30.2 | 15 | USGS Water Data Vermilion Bay - 07387040 |
| 8 | Sister Lake | LA | 29.2491 | -90.9211 | 3.0 | 18.0 | 13.2 | 30.7 | 16 | USGS Water Data Sister Lake - 07381349 |
| 9 | Barataria Bay | LA | 29.3985 | -90.0411 | 2.2 | 17.0 | 13.6 | 30.6 | 15 | USGS Water Data Culch Plant/Hackberry Bay - 073802512 |
| 10 | Lake Fortuna | LA | 29.6335 | -89.5636 | 1.7 | 18.0 | 12.6 | 30.3 | 16 | USGS Water Data Lake Fortuna - 07374526 |
| 11 | Grand Bay | AL | 30.3486 | -88.4185 | 14.0 | 28.8 | 12.8 | 30.6 | 19 | NERR Centralized Data. Grand Bay - Point Aux Chenes Bay GNDPCWQ |
| 12 | Saint Vincent Sound | FL | 29.6747 | -85.0583 | 9.7 | 31.6 | 13.9 | 30.3 | 23 | NERR Centralized Data. Apalachicola Bay - Dry Bar APADBWQ |
| 13 | East Cove | FL | 29.7021 | -84.8802 | 10.1 | 31.2 | 14.1 | 30.4 | 23 | NERR Centralized Data. Apalachicola Bay - Cat Point APACPWQ |
| 14 | Timucuan Preserve | FL | 30.4411 | -81.4390 | 27.8 | 35.9 | 14.5 | 29.2 | 20 | National Parks Service Continuous Water Data - Timucuan Preserve |
| 15 | Sapelo Island | GA | 31.4179 | -81.2960 | 20.8 | 32.2 | 12.4 | 29.7 | 25 | NERR Centralized Data. Sapelo Island - Lower Duplin SAPLDWQ |
| 16 | Doboy Sound | GA | 31.4533 | -81.3627 | 17.0 | 31.0 | 12.8 | 30.0 | 16 | USGS Water Data Meridian Landing - 022035975 |
| 17 | Skidaway River | GA | 31.9863 | -81.0052 | 21.5 | 29.8 | 11.0 | 30.0 | 3 | Biological and Chemical Oceanography Data Management Office |
| 18 | Bull River | GA | 31.0141 | -80.8841 | 16.8 | 28.6 | 11.8 | 39.4 | 18 | National Parks Service Continuous Water Data - Fort Pulaski NM - Lazaretto Creek Dock |
| 19 | Charleston Harbor | SC | 32.7804 | -79.9236 | 22.0 | 29.0 | 12.0 | 29.4 | 16 | USGS Water Data Charleston Harbor - 021720710 |
| 20 | Murrells Inlet | SC | 33.5240 | -79.0619 | 32.2 | 35.6 | 10.8 | 28.3 | 15 | BCCWMS Coastal Volunteer Monitoring Data |
| 21 | Cape Fear River | NC | 33.9547 | -77.9350 | 13.2 | 29.3 | 10.0 | 29.4 | 23 | NERR Centralized Data. North Carolina - Zeke's Basin/Cape Fear River NOCZBWQ |
| 22 | Masonboro Island | NC | 34.1400 | -77.8625 | 21.7 | 34.4 | 10.8 | 28.7 | 6 | UNCW's Shellfish Research Hatchery at the Center for Marine Science and NCNERR - CMSDOCK |
| 23 | Pamlico Sound | NC | - | - | - | - | - | - | - | - |
| 24 | Wreck Shoal | VA | - | - | - | - | - | - | - | - |
| 25 | Deep Water Shoal | VA | - | - | - | - | - | - | - | - |
| 26 | Eastern Shore | VA | 37.6076 | -75.6858 | 27.3 | 32.3 | 6.1 | 27.8 | 7 | VIMS Eastern Shore Laboratory: Custis Channel/Wachapreague |
| 27 | Potomac River | MD | 38.3626 | -76.9905 | 2.7 | 12.0 | 4.7 | 27 | 37 | Maryland DNR/Eyes on the Bay - Lower Potomac River, Morgantown Bridge |
| 28 | Cape Shore | NJ | 39.0850 | -75.1860 | 17.4 | 26.7 | 7.6 | 24.9 | 18 | Delaware Water Quality - CEMA at University of Delaware |
| 29 | Hope Creek | NJ | 39.4550 | -75.5600 | 1.3 | 10.0 | 8.1 | 26.7 | 18 | Delaware Water Quality - CEMA at University of Delaware |
| 30 | Ash Creek | CT | 41.1884 | -73.1212 | 1.1 | 27.0 | 4.5 | 24.4 | 4 | USGS Water Data - Housatonic River NR Nells Island NR Stratford, CT - 01208837 |

| | | | | | | | | | | |
|---|---|---|---|---|---|---|---|---|---|---|
| 31 | Long Island Sound | CT | - | - | - | - | - | - | - | - |
| 32 | Narrow River | RI | - | - | - | - | - | - | - | - |
| 33 | Narragansett Bay | RI | 41.5783 | -71.3211 | 29.2 | 31.7 | 3.5 | 21.0 | 22 | NERR Centralized Data. Narragansett Bay T-Wharf Bottom NARTBWQ |
| 34 | Plum Island Sound | MA | 42.7625 | -70.8577 | 11.8 | 30.8 | 11.6 | 24.7 | 7 | PIELTER Long-Term Monitoring Data |
| 35 | Great Bay | NH | 43.0524 | -70.9118 | 5.9 | 28.6 | 7.2 | 24.2 | 27 | NERR Centralized Data. Great Bay - Squamscott River GRBSQWQ |
| 36 | Orrs Cove | ME | 43.9860 | -69.5500 | 26.2 | 31.4 | 7.5 | 21.5 | 7 | University of Maine EPSCOR |
| 37 | Damariscotta River | ME | 43.8645 | -69.8976 | 30.5 | 32.0 | 8.3 | 19.1 | 4 | University of Maine EPSCOR |
| 38 | Salt Bay | CN | 43.7907 | -65.8361 | 21.3 | 30.5 | 1.8 | 20.3 | 3 | Nova Scotia Gov Water Quality Data |
| 39 | Eel Lake | CN | - | - | - | - | - | - | - | - |
| 40 | Prince Edward Island | CN | - | - | - | - | - | - | - | - |

*Supp. Table S3: Mutation table with all SNPs and significance values from all analyses*

Supp. Table S3 mutation table

*Supp. Table S4: lfmm2 global high temperature enrichment analysis*

Supp. Table S4 global high temp

*Supp. Table S5: lfmm2 global low temperature enrichment analysis*

Supp. Table S5 global low temp

*Supp. Table S6: lfmm2 Atlantic high salinity enrichment analysis*

Supp. Table S6 Atlantic high salinity

*Supp. Table S7: lfmm2 Atlantic low salinity enrichment analysis*

Supp. Table S7 Atlantic low salinity

*Supp. Table S8: lfmm2 Gulf high salinity enrichment analysis*

Supp. Table S8 Gulf high salinity

*Supp. Table S9: lfmm2 Gulf low salinity enrichment analysis*

Supp. Table S9 Gulf low salinity

*Supp. Table S10: pcadapt gene ontology enrichment analysis*

Supp. Table S10 pcadapt

*Supp. Table S11: OutFLANK gene ontology enrichment analysis*

Supp. Table S11 OutFLANK

*Supp. Table S12: Top 100 consensus genes*

Supp. Table S12 Top 100 consensus genes

<u>*Supplemental Figures*</u>

*Fig. S1. Global snmf cross-entropy plot*

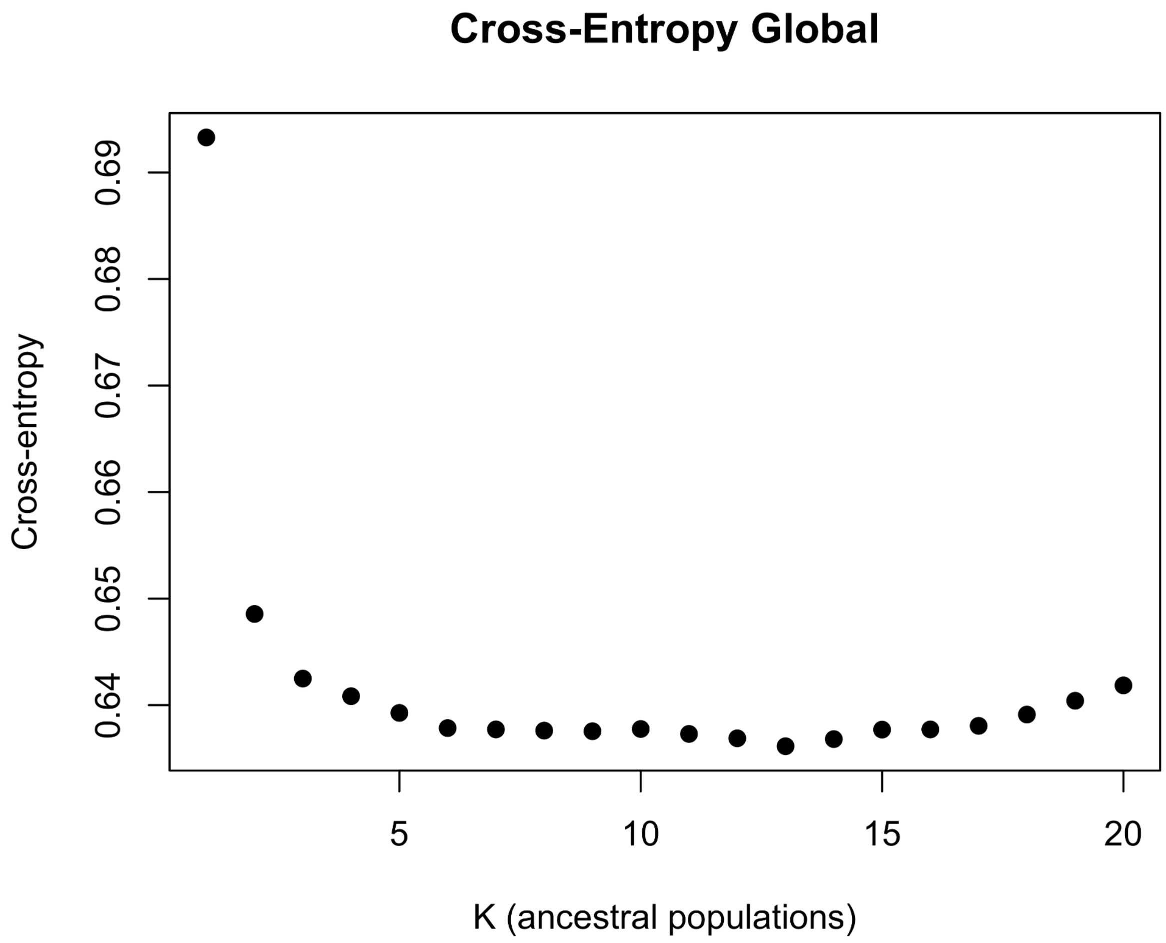


***Figure S1.*** Cross-entropy values from SNMF ancestry analysis for the global dataset. The selected cross-entropy value was K=2 (0.6481).

*Fig. S2. Gulf snmf cross-entropy plot*

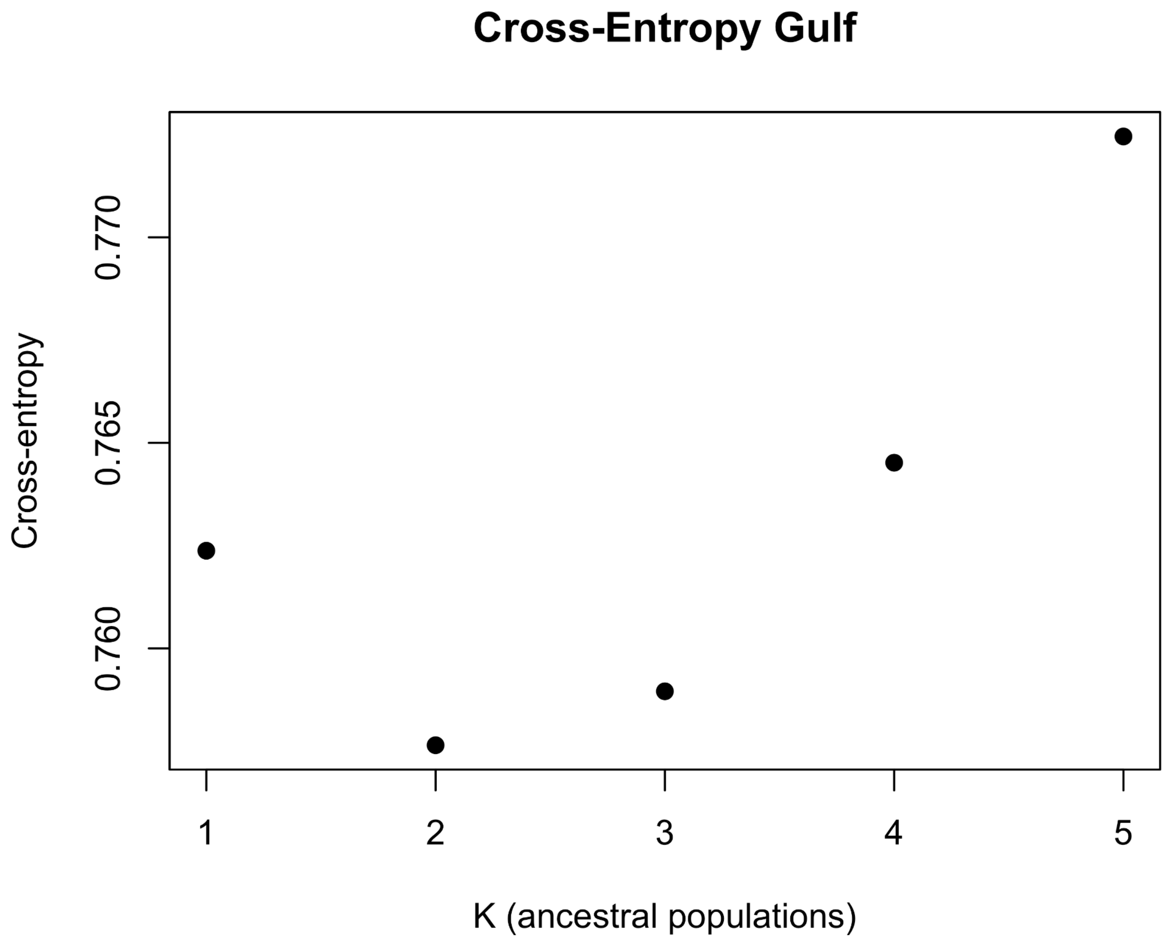


***Figure S2.*** Cross-entropy values from SNMF ancestry analysis for the Gulf dataset. The lowest cross-entropy value was K=2 (0.7576).

*Fig. S3. Atlantic snmf cross-entropy plot*

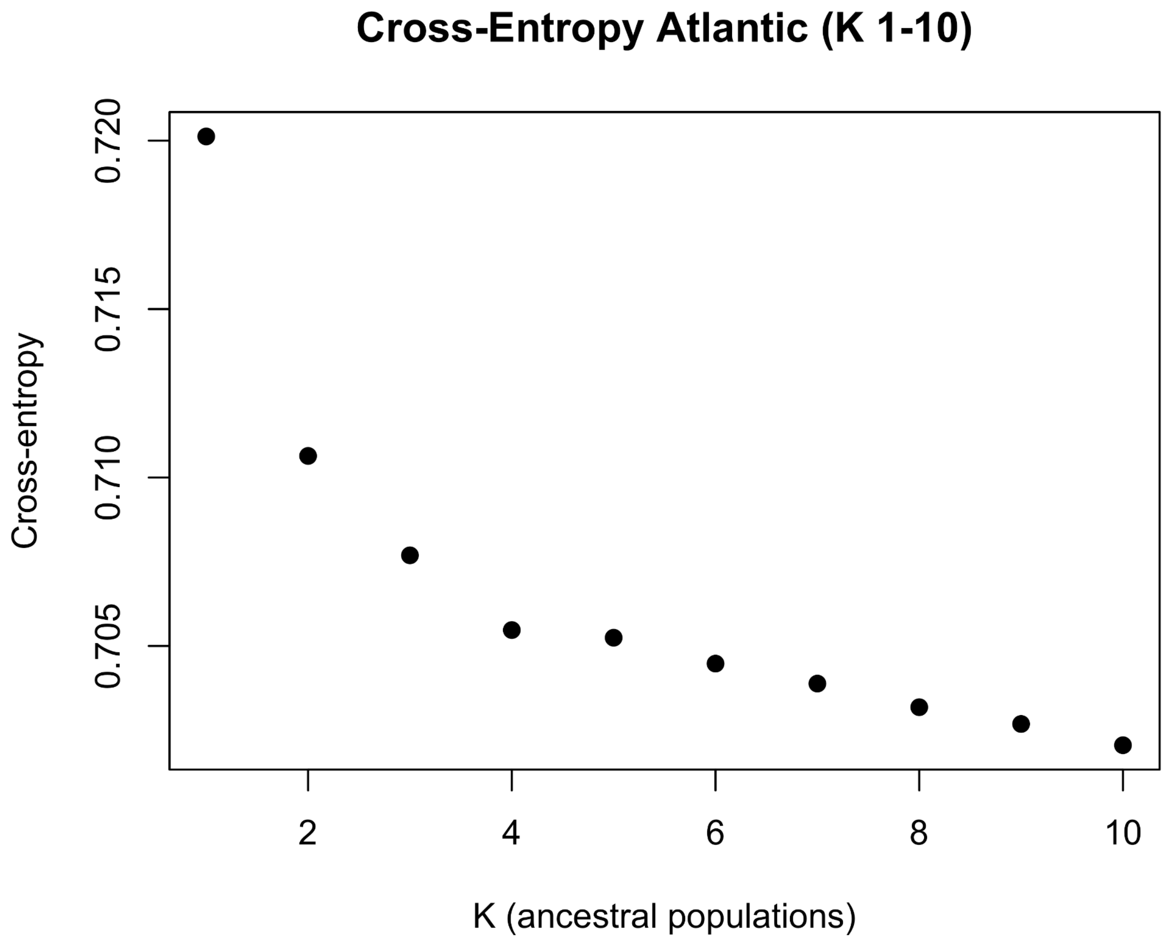


***Figure S3.*** Cross-entropy values from SNMF ancestry analysis for the Atlantic dataset. The chosen cross-entropy value was K=5 (0.7052).

*Fig. S4. Local PCA MDS1 window plot for all chromosomes*

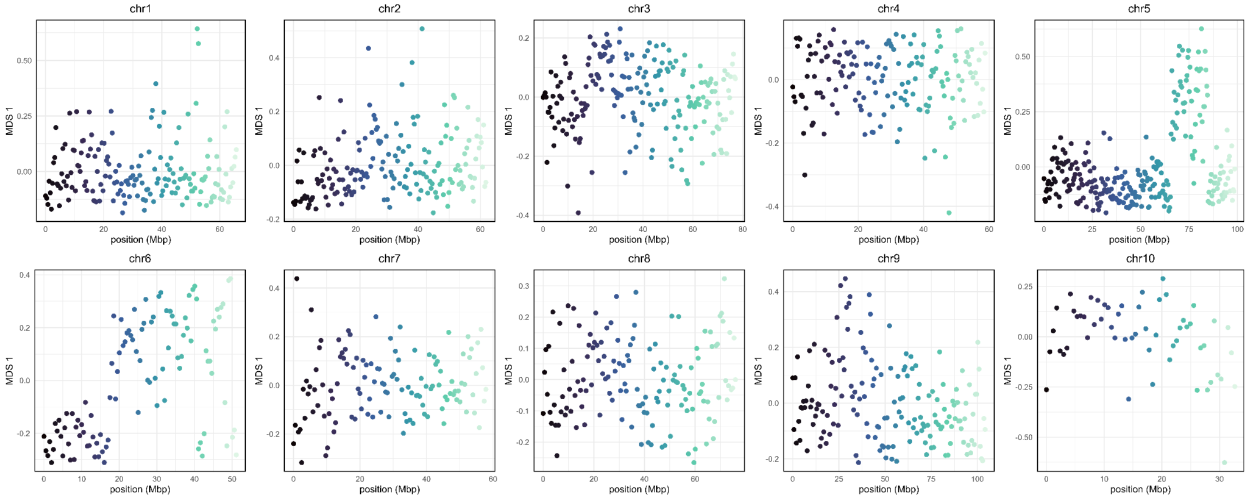


***Figure S4.*** Chromosomal regions for detection of structural variants from local PCA analysis. MDS1 is plotted by SNP position (Mbp, x-axis) on each chromosome. Colors correspond to the position of the window position within the chromosome, where the darkest shade starts at 0 Mbp. Outlier SNP windows are visualized as areas of divergent positive or negative MDS1 values.

*Fig. S5. Synteny plot with chromosome-linkage group misalignments*

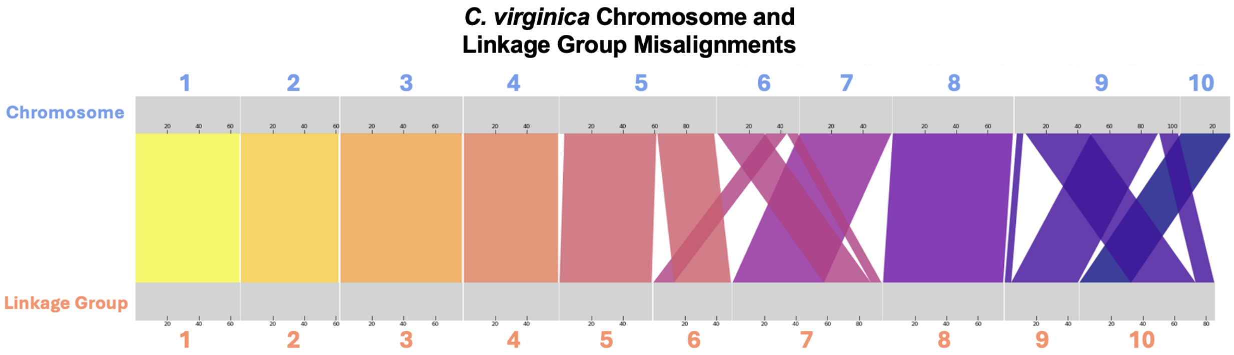


***Figure S5.*** Synteny plot with chromosome (top) and linkage group (bottom) misalignments in the *C. virginica* genome assembly.
